# Cytoplasmic Switch of ARS2 Isoforms Promotes Nonsense-Mediated mRNA Decay and Arsenic Sensitivity

**DOI:** 10.1101/2021.07.08.451683

**Authors:** Monica Mesa-Perez, Phineas T. Hamilton, Alex Miranda-Rodriguez, Nicholas Brodie, Connor O’Sullivan, Jennifer Christie, Bridget C. Ryan, Robert L. Chow, David Goodlett, Christopher J. Nelson, Perry L. Howard

## Abstract

The life of RNA polymerase II (RNAPII) transcripts is shaped by the dynamic formation of mutually exclusive ribonucleoprotein complexes (RNPs) that direct transcript biogenesis and turnover. A key regulator of RNA metabolism in the nucleus is the scaffold protein ARS2 (arsenic resistance protein 2), bound to the cap binding complex (CBC). We report here that alternative splicing of *ARS2’s* intron 5, generates cytoplasmic isoforms that lack 270 amino acids from the N-terminal of the protein and are functionally distinct from nuclear ARS2. Switching of ARS2 isoforms within the CBC in the cytoplasm has dramatic functional consequences, changing ARS2 from a NMD inhibitor to a NMD promoter that enhances the binding of UPF1 to NCBP1, ERF1 and DHX34, favouring SURF complex formation, SMG7 recruitment and transcript degradation. ARS2 isoform exchange is also relevant during arsenic stress, where cytoplasmic ARS2 promotes a global response to arsenic in a CBC-independent manner. We propose that ARS2 isoform switching promotes the proper recruitment of RNP complexes during NMD and the cellular response to arsenic stress. The existence of non-redundant ARS2 isoforms is relevant for cell homeostasis, stress response, and cancer treatment.

## Introduction

The life of an mRNA is highly coordinated from its beginnings during transcriptional initiation to its inevitable degradation. In the nucleus, the coordination is guided in part by the nuclear cap binding complex (CBC), consisting of NCBP1 and NCBP2, bound to the 7-methyl guanosine cap of RNAPII transcripts^1^. ARS2/SRRT plays an important scaffolding role within the CBC by helping to synchronize the dynamic assembly and disassembly of mutually exclusive complexes that regulate mRNA splicing, degradation, and export^1–7^. CBC-ARS2 bound mRNAs are exported to the cytoplasm, where NCBP1 is required for the pioneering round of translation and for promoting transcript degradation through nonsense mediated decay (NMD)^8,9^. Although ARS2 is translocated to the cytoplasm with the CBC, it is unknown whether ARS2 participates in CBC-related functions in the cytoplasm.

NMD is a translation-dependent surveillance process that promotes the cytoplasmic degradation of mRNA with premature termination codons (PTC), to prevent the synthesis of dysfunctional proteins and maintain mRNA homeostasis^10,11^. During nuclear splicing, the multiprotein Exon Junction Complex (EJC) is deposited onto the mRNAs 24 nt upstream of the spliced junctions^12^. EJCs are exported to the cytoplasm with the mRNAs and are dissociated by the ribosome during translation. By recruiting several factors involved in splicing, transport, translation and NMD, the EJC establishes a molecular link between nuclear splicing and cytoplasmic NMD^13–15^. Mammalian-cell NMD generally occurs when translation terminates more than 50–55 nucleotides upstream of an EJC complex^16^. In this scenario, the termination codon is recognized as a PTC, which triggers RNP recruitment and remodelling. First, UPF1 is recruited to the stalled ribosome by ERF1, ERF3 and NCBP1. Subsequently, SMG1 kinase joins the complex to generate the SURF complex (SMG1–UPF1– ERF1–ERF3). Next, in a step dependent on DHX34 and NCBP1, UPF1 interacts with the downstream UPF2/3 within the EJC to form the decay-inducing (DECID) complex, triggering SMG1 activation. The phosphorylation of UPF1 by SMG1 leads to translation inhibition and promotes mRNA degradation through the recruitment of SMG6, SMG5–SMG7 and mRNA decay factors^15^. Importantly, inhibition of the NMD downstream of the DECID complex results in UPF1 hyperphosphorylation^17^. Previous work has shown that CBC complex component NCBP1 promotes NMD in the cytoplasm^9,18^. Additional studies have reported the interaction between ARS2 and several components of the NMD pathway^5,19,20^. However, the question of whether ARS2 participates in NMD has not been addressed.

Another unanswered question of ARS2 function relates to its known role in the cellular response to arsenic. Arsenic is not only a widespread environmental contaminant, but is also used to treat certain cancers, such as acute promyelocytic leukemia (APL)^21^. The initial study of *Ars2* showed that overexpression conferred resistance to arsenic^22^. The cDNA used in this study encoded only a small portion of the extreme C-terminus of full-length ARS2 and generated a dominant negative. It was subsequently found through knockdown studies that full length *Ars2* conferred arsenic sensitivity^3^. However, we still do not know how ARS2’s role in RNA metabolism relates to the cellular response to arsenic.

In this study, we demonstrated that alternative splicing of intron 5 generates cytoplasmic isoforms of ARS2 (herein named ARS2c), that lack 270 amino acids from the N-terminal of the protein. Despite sharing the middle, C-terminal domains and unstructured regions (including the RNA and CBC binding regions) with nuclear ARS2 (ARS2n), ARS2c is functionally distinct from ARS2n. The differential biological roles of ARS2 isoforms is evidenced in the regulation of NMD, where our data shows that both ARS2n and ARS2c work in tandem to modulate the pathway. ARS2n primarily functions in the nucleus with the CBC to regulate mRNA splicing, degradation, and export^1–7^. We propose that, once the mRNPs enter the cytoplasm, an isoform switch occurs in which ARS2c replaces ARS2n within the CBC. The association of ARS2c with the CBC increases the interaction between UPF1 and ERF1, NCBP1 and DHX34, promoting the formation of the SURF complex at the PTC, SMG7 recruitment and transcript degradation though NMD. Regulation of the NMD pathway by ARS2 isoforms seems to be relevant *in vivo*, since ARS2 expression in tumours is associated with a strong enrichment of NMD across solid tumours evaluated from the Cancer Atlas (TCGA). The differential biological roles of ARS2 isoforms is also seen during stress response. We show that ARS2c and not ARS2n is specifically induced during arsenic stress and it is responsible for arsenic sensitivity. Knock down of ARS2c is sufficient to abrogate proteomic remodeling and cellular response to arsenic, in a process that it is independent of the CBC complex. ARS2c mediated arsenic sensitivity is, to our knowledge, the first described function for mammalian ARS2 that is potentially independent from the CBC complex.

The role of ARS2 isoforms in the regulation of NMD and arsenic sensitivity is likely to be relevant in cancer. Arsenic trioxide has been proposed as a potential treatment for pancreatic cancer^23^, while missregulation of NMD, due to UPF1 mutation, is a molecular signature in pancreatic adenosquamous carcinoma^24^. Our study shows that ARS2 expression correlates with better survival in pancreatic cancer, but inferior survival in several other cancer types. Given the key role of ARS2 isoforms in NMD and arsenic sensitivity, understanding the functions of ARS2 isoforms in cancer is timely.

## Materials and Methods

### Cell Culture, transfection, and generation of stable cell lines

C_2_C_12_ (ATCC: CRL-1772), HeLa (ATCC: CCL-2), HEK 293T (ATCC: CRL-11268) or Flp-In T-Rex (Invitrogen: R78007) cell lines were cultured in Dulbecco’s Modified Eagle’s Medium High Glucose (DMEM-high glucose, Hyclone) and supplemented with 10% fetal bovine serum (FBS, Hyclone) at 37°C in 5% CO_2_ buffered incubators. To generate stable cell lines expressing Biotin ligase-ARS2n/ARS2c1/ARS2c2 or control, Flp-In T-REx cells were transfected with a 9:1 ratio of pOG44 (Thermo Fisher) to pcDNA5 integration vector and allowed to recover for 48h prior to selection with hygromycin (InvivoGen). Stable clones were pooled and tested for expression with the addition of 1ug/ml tetracycline (Sigma) for 24h. Transfections were performed using JetPrime reagent (VWR) as per the manufacturer’s instructions. Small interfering RNAs (siRNAs) were purchased from Integrated DNA Technologies (IDT) and AllStars negative-control siRNA (Qiagen) was used as a control siRNA. shRNAs were purchased from the HuSH™ shRNA collection at Origene in the vector pGFP-V-RS and scramble control was used as control. All siRNA/shRNA sequences are reported in Supplemental table 1.

### Plasmids

Expression vectors for stable integration and tetracycline-inducible expression, were generated by subcloning synthetized ARS2n/ARS2c1/ARS2c2-3xFLAG (GenScript) into a modified pcDNA5/FRT/TO vector containing BirA, kindly provided by C. Nelson^25^. BIOD2-ARS2n/ARS2c1/ARS2c2-3xFLAG vectors were generated by synthetizing BioID2 and substituting BirA on the previous constructs (GenScript). BIOD2-nlsKO-ARS2n mutant was generated by mutagenesis of BIOD2-ARS2n and subcloned in pcDNA3.1(+)-N-eGFP (GenScript). All constructs were validated by sequencing. Enhanced GFP-C1 (eGFP-C1) was used as a green fluorescent protein (GFP) expression control, eGFP-ARS2n and eGFP-ARS2c1 were generated as described previously ^26^. The Firefly luciferase 5xbox b reporter (FLuc-5Xboxb) and firefly luciferase control (FLuc) were generated by subcloning FLuc-5Xbox b from plasmid pAc5.1C-Fluc-STOP-5boxb (Addgene) into pcDNA3.1(+) (Life Technologies). Renilla luciferase control plasmid (RLuc) was obtained from Promega. λN was synthetized and cloned into pcDNA3.1 (-) ARS2^27^ or pcDNA3.1 (-) alone generating ARS2-λN and λN control respectively. λN-RNPS1 plasmid was generated by synthesis of RNPS1 and cloning on λN pcDNA3.1 (-) plasmid. pNMD+ and pNMD-reporter plasmids were kindly provided by K. Lukyanov^28^. RNT1-GFP was a gift from H. Dietz (Addgene plasmid # 17708)^29^. pAc5.1C-FLuc-Stop-5BoxB was a gift from Elisa Izaurralde (Addgene plasmid # 21301)^30^. CBP20-3flag was a gift from Torben Heick Jensen and John LaCava^31^.

### RNA isolation, cDNA generation, quantitative real-time PCR and real-time PCR

To guarantee data robustness and consistency, three individually treated wells from a 6 well plate were pooled into 1 RNA column. This RNA column was defined as “biological replicate”. Three technical replicates were run per biological replicate. For every analyzed gene, 3 to 6 biological replicates were included, leading to 9-18 datapoints per condition that represent 9-18 individually treated wells. Total RNA was isolated using RNeasy plus kit (Qiagen). To guarantee extensive elimination of genomic DNA, RNA samples were additionally treated with DNAseI (Thermofisher). 1ug of total RNA was reverse transcribed using High-Capacity cDNA Reverse Transcription kit (Thermofisher). PCR was performed using Q5 High-Fidelity DNA polymerase (NEB) while qPCR products were amplified using Ssofast EvaGreen Supermix (BioRad) on a Stratagene MX3000P qPCR system. All primer sequences are reported in Supplemental table 1.

### Confocal microscopy

HEK 293T cells were seeded on glass coverslips (neuVitro) and transfected with BIOD2-ARS2n/ARS2c1/ARS2c2-3xFLAG or control during 48 hours in complete DMEM media. Cells were washed with PBS and fixed for 15 minutes with 4% paraformaldehyde in PBS at RT. Next, cells were permeabilized in 0.25% Triton X-100 for 15 minutes at RT and blocked with 1% BSA in PBS/Tween (0.01%), 1 hour at RT. Primary (anti-flag mouse 1:200, Cell Signalling) and secondary antibody (anti-mouse Alexa 488 1:500, Thermofisher) were diluted in blocking buffer solution and incubated overnight at 4^°C^ or 1 hour at RT, respectively. Hoescht 3334 (1:2000) was added at RT for 5 minutes to label nuclei. Images were acquired using a confocal microscope (Nikon) and processed in ImageJ.

### Western Blot

Cell lysates, quantified by Pierce BCA assay kit (Thermofisher) and resuspended in Laemmli sample buffer, were resolved by SDS-PAGE and transferred to polyvinylidene difluoride (PVDF) membranes. Membranes were blocked in Intercept (TBS) Blocking Buffer (Licor) for 1 hour at 37°C. Primary and secondary antibodies were diluted in Blocking buffer/Tween (0.1%) and membranes were incubated overnight at 4°C or 1 hour at RT, respectively. Images were revealed and analyzed using Odyssey CLx (Licor) and Image Studio Lite software. Total proteins were detected using Revert 700 Total protein Stain kit (Licor).

### Antibodies

The following antibodies were used in this study: ARS2 (XL14.1, 1:2000) generously provided by the Ludwig Institute for Cancer Research, Flag (1:1000, Cell signalling), Actin (1:4000, Sigma), TBP (1:1000, Cell Signalling), Tubulin (1:1000, Cell Signalling), GFP (1:1000, Cell Signalling), eRF1 (1:1000, Thermofisher), eIF3E (1:1000, Abclonal), eIF3F (1:1000, Abclonal), eIF3K (1:1000, Abclonal), Magoh (1:1000, Abclonal), MagohB (1:1000, Abclonal), eIF4A3 (1:1000, Abclonal), SMG7 (1:1000, Thermofisher), SMG1 (1:500, Santa Cruz), DHX34 (1:1000, Cedarlane), Streptavidin 680 (1:15000, Invitrogen), UPF1 (1:500, Santa Cruz), UPF1 (1:1000, Cell Signalling), NCBP1 (1:1000, Cell Signalling), Phospho-(Ser/Thr) ATM/ATR Substrate (1:1000, Cell Signalling).

### Protein fractionation

C_2_C_12_ cells were washed with buffer H (20mM HEPES pH:8, 2mM MgCl_2_, 0.1mM EGTA, 1mM EDTA) and lysed for 10 minutes at 4°C in buffer I (2x buffer H, 0.2% NP-20, protease inhibitor cocktail). Cell’s nuclei were pelleted at 16000 ⨯ g for 10 minutes and supernatant was stored as cytoplasmic fraction. Cold RIPA buffer with protease inhibitor cocktail was added to the pelleted nuclei and samples were incubated 30 minutes on ice. Cell debris was pelleted at 16000 x g for 20 minutes and supernatant was stored as nuclear fraction. Nuclear and cytoplasmic fractions were quantified by BCA Protein Assay kit (Thermofisher).

### Affinity capture of biotinylated proteins

Purification of biotinylated proteins was performed as outlined by^32^. Flp-In T-REx cells stably expressing tetracycline inducible ARS2n-BirA/control or BioID2-ARS2n/ARS2c1/ARS2c2/control, were treated for 24h with 10-200ng/ml of tetracycline. HEK 293T cells were transfected with BIOD2-ARS2n/ARS2c1/ARS2c2/-3xFLAG or control and incubated for 24h. All the conditions were incubated an additional 24h in complete DMEM media supplemented with 50uM of biotin (Sigma). Five individually treated 10 cm^2^ dishes, corresponding to around 50⨯10^6^ total cells were pooled together and considered 1 biological replicate. One biological replicate (5⨯10cm^2^ dishes) was used for tetracycline inducible BioID2-ARS2n/ARS2c1/ARS2c2/control, and three biological replicates (15⨯10cm^2^ dishes) were used for the rest of the evaluated conditions (tetracycline inducible ARS2n-BirA/control and BIOD2-ARS2n/ARS2c1/ARS2c2/-3xFLAG/control overexpression). Cells were washed three times in PBS and lysed directly on the plate with 500ul Lysis Buffer (50 mM Tris pH 7.4, 500 mM NaCl, 0.2% SDS, 1 mM DTT, and protease inhibitors cocktail). Triton X-100 was added to a final volume of 2%. Lysates were sonicated on 30% amplitude for 2 cycles of 30 seconds with 2 minutes rest, using a sonic dismembrator (Fisher Scientific). Lysates were diluted in chilled 50mMTris pH7.4 and subjected to a final round of sonication. Insoluble cellular debris was cleared by centrifugation at 16 000 x g for 20 minutes at 4°C. Cleared extracts were incubated with 200 ul of Dynabeads MyOne Streptavidin C1 (Thermo Fisher) and incubated overnight at 4°C. The following morning, beads were washed two times with Wash Buffer 1 (2% SDS), once with Wash Buffer 2 (0.1% deoxycholic acid, 500 mM NaCl, 1 mM EDTA, 50 mM HEPES ph7.5 and 1% Triton X-100), once in Wash Buffer 3 (10 mM Tris pH 7.4, 250 mM LiCl, 0.5% NP-40, 0.5% deoxycholic acid, and 1mM EDTA), and finally two times with Wash Buffer 4 (50 mM Tris pH 7.4 and 50 mM NaCl), pelleting beads using a magnetic rack. For western blotting, beads were resuspended in Laemmli sample buffer supplemented with 50 uM biotin, incubated for 10 minutes, and boiled for 10 minutes before loading on an SDS-PAGE gel. Total protein was detected using Silver Stain kit (Thermofisher) or Coomasie stain, and BIOD2-ARS2n/ARS2c1/ARS2c2/control concentration was estimated by extrapolation on a BSA curve ran on the same gel. Equal amounts of proteins were used as input for LC-MS/MS.

### Protein identification by mass spectrometry

BioID samples were processed for mass spectrometry as outlined by^25^. Briefly, streptavidin beads were washed with 50 mM NH4HCO3, resuspended in 50 mM NH_4_HCO_3_ containing 5 mM dithiothreitol, and heated at 75°C for 10 minutes. Iodoacetamide was added to a final concentration of 10 mM to each sample followed by incubation in the dark at RT for 1 hour. Afterwards, 1 mM CaCl_2_ and 1 ug of sequence-grade trypsin (Promega) were added and incubation continued overnight. The following morning, trifluoroacetic acid (TFA) was added to a final concentration of 0.5% (v/v). Beads were pelleted using a magnetic rack and the supernatant removed. A second elution of digested peptides with 0.5% TFA was performed and the supernatant pooled. Digested peptides were passed over ZipTips (Millipore) and eluted with 0.5% formic acid/ 80% acetonitrile. Samples were diluted ½ in water, lyophilized and store at -80°C until use.

### Mass spectrometry acquisition using Orbitrap Fusion Tribrid mass spectrometer (Thermo Scientific)

Digested samples (6 µL) were separated by on-line reverse phase chromatography using a Thermo Scientific EASY-nLC 1000 system with a reverse-phase pre-column Magic C18-AQ (100 µm I.D., 2.5 cm length, 5 µm, 100 Å) and an in-house prepared reverse phase nano-analytical column Magic C-18AQ (75 µm I.D., 15 cm length, 5 µm, 100 Å, Michrom BioResources Inc, Auburn, CA), at a flow rate of 300 nl/min. The chromatography system was coupled on-line with an Orbitrap Fusion Tribrid mass spectrometer (Thermo Fisher Scientific, San Jose, CA) equipped with a Nanospray Flex NG source (Thermo Fisher Scientific). Solvents were A: 2% Acetonitrile, 0.1% Formic acid; B: 90% Acetonitrile, 0.1% Formic acid. After a 348 bar (∼ 3 µL) pre-column equilibration and 348 bar (∼ 3 µL) nanocolumn equilibration, samples were separated by a 60-minute gradient (0 min: 0%B; 52 min: 45%B; 2 min: 100%B; hold 6 min: 100%B). Data-dependent acquisition Orbitrap survey spectra were scheduled at least every 3 seconds, with the software determining “Automatic” number of MS/MS acquisitions during this period^33^.

### MS data analysis

Raw files were created by XCalibur 4.3.73.11 (Thermo Scientific) software and analyzed with PEAKS Client 7.0 software suite (Bioinformatics Solutions Inc.). Database search parameters as follows: precursor tolerance 5 ppm; MS/MS tolerance 0.035 Da; Trypsin enzyme 2 missed cleavages; Orbi-TRAP instrument type; fixed modification: Carbamidomethylation (C); variable modifications: Oxidation (M), Acetylation (K, N-term) Biotinylation. SwissProt_20200305 Database (561911Sequences /202173710 residues)^34^. To identify significant interactors with the different ARS2 isoforms, we tested for differences in unique peptide counts across different baits using triplicate replicates of each bait condition, as well as additional negative controls from^35,36^. Unique peptide counts were used as input for SAINTexpress (v3.6.3; https://www.sciencedirect.com/science/article/abs/pii/S1874391913005381). For each bait, significant prey were defined as those with a SAINT score > 0.7 and > 2 average unique peptides across conditions. Over-representation analysis of prey for each bait was conducted using the enricher function of ClusterProfiler (https://pubmed.ncbi.nlm.nih.gov/22455463/). Data visualizations were generated in R (v > 4.0), using the ComplexHeatmap (https://academic.oup.com/bioinformatics/article/32/18/2847/1743594) package. Functional annotation maps of ARS2 isoform interactomes were generated by mining for enriched GO ‘‘Biological Process’’ terms using the CluoGo plugin^37^ within the Cytoscape framework^38^. ClueGO is a user-friendly plugin that allows the decoding and visualization of functionally grouped GO terms in the form of networks. The size of the nodes shows the term significance after Bonferroni correction. Only GO terms with a P-value < 0.01 were considered significant. A kappa score was calculated reflecting the relationships between the terms based on the similarity of their associated genes, which was set to 0.5 as the threshold in this study. The Organic algorithm that determines node positions based on their connectivity was used for laying out the networks. Protein-Protein interaction networks were generated using STRING^39^. Saint analyses are reported in Supplemental tables 2-4. Mass spectrometry proteomics data have been deposited to the ProteomeXchange Consortium via the PRIDE partner repository.

### Protein Immunoprecipitation

**Co-IP:** Cells co-transfected with BIOD2-ARS2n/ARS2c1/ARS2c2/BioID2-3xFLAG vectors and GFP-UPF1 or GFP control were lysed in RIPA lysis for 30 minutes at 4°C. Cell debris was removed by centrifugation (20 minutes at 16 000 x g). Extracts were quantified by Pierce BCA Protein quantification kit (Thermofisher) and incubated with anti-FLAG beads or GFP Trap magnetic beads (Chromotek) overnight at 4°C. Beads were washed three times with lysis buffer-PBS at 1:4 and eluted with 3xFLAG peptide (Sigma-Aldrich) or Laemmli sample buffer, respectively. **Endogenous IP:** Cell lysates were obtained as before. Cell extracts were incubated overnight at 4°C, with 1 ug of anti-UPF1 or anti-ARS2 antibodies and protein A/G magnetic beads (Genscript). The next day, beads were washed with PBS and resuspended in Laemmli sample buffer. All samples were heated at 95°C for 10 minutes.

### Cell survival assays: WST1 and Crystal Violet

C_2_C_12_ cells were transfected with *Ars2c*/*Ars2*(all)/*Ncpb1*/*Ars2c*+*Ncbp1* or control RNAi for 24 hours. Media was changed to Arsenic (35 uM/40 uM) and cells were incubated for 24, 48, 72, 96, or 120 hours after the arsenic treatment. Cells transfected with the described siRNA for 48 hours but untreated with arsenic were defined as point zero. Arsenic media was changed every 3 days. For WST1 assays, WST1 solution (Sigma) was added to the supernatant of the cells and incubated for 4 hours at 37°C in 5% CO_2_ buffered incubator. Absorbance was measured at 450 nm on BioTek Epoch 2 microplate spectrophotometer, normalizing samples against a blank well. For crystal violet assays, cells were washed with PBS and fixed in 10% formalin for 10 minutes at RT. After washing the plates with water, crystal violet solution was added, and plates were incubated on a rocker for 20 minutes at RT. Crystal violet solution was removed with several washes of water and plates were dried 24 hours at RT. After taking pictures, 10% acetic acid was added to the plates and absorbance was measured at 570 nm on BioTek Epoch 2 microplate spectrophotometer, normalizing samples against a blank well.

### NMD tethering assay

HeLa cells were transfected with either control siRNA or *UPF1*si for two consecutive days, and on day 3 were co-transfected with NMD-firefly (FLuc-5xboxb) and Renilla luciferase (RLuc) and either λN or λN-ARS2. Firefly and Renilla luminescence were analyzed 72h after the first transfection using the Dual-Glo Luciferase Assay System (Promega) and a Perkin Elmer Victor3V 1420 multi-label plate reader. Firefly luminescence was normalized to Renilla luminescence.

### NMD reporter assay

HeLa and C_2_C_12_ cells were transfected with *ARS2*si/*ARS2c*si/*UPF1*si or control. Media was changed 16 hours post-transfection and each condition was transfected again with either pNMD+ or pNMD-vectors. Flow cytometry and RT-qPCR analysis were performed 24 or 48h after the second transfection. For flow cytometry 1.0×10^5^ events were acquired using a BD FACSCalibur. Cells expressing either Katushka (pTurboFP635-N vector, Evrogen) or TagGFP2 (pTagGFP2-N vector, Evrogen) were used as controls for the crosstalk of the TagGFP2 signal into the red channel and the Katushka signal into the green channel. Data analysis was performed as described by the developers of the assay^40^.

### Immunofluorescence

Cells were transfected with the indicated GFP-ARS2n/nlsKO-ARS2n/control constructs. Twenty-four hours post-transfection, cells were live imaged using a 20x objective on a Leica DMIRE2 inverted fluorescence microscope. Hoescht 3334 (1:2000) was added at RT for 5 minutes to label nuclei. Images were processed in ImageJ.

### TCGA bioinformatic analysis

To examine correlates of ARS2 expression in cancer samples, we accessed the RNA-sequencing and clinical outcomes data for cases profiled by the Cancer Genome Atlas (TCGA), using recently harmonized pan-cancer data (https://gdc.cancer.gov/about-data/publications/pancanatlas). We retained cases present in both outcomes and mRNA expression datasets (for expression analyses). For survival analyses, we removed cases with missing survival data, or cancer types with fewer than 30 evaluable events from consideration. Expression data were log2(x + 1) transformed prior to further analysis.

We estimated log hazard ratios for *SRRT* (ARS2) expression in each cancer using univariable Cox proportional hazards models, with log *SRRT* expression as the predictor variable. P values for *SRRT* effects were multiple-test corrected using the Benjamini Hochberg method. To compute gene set enrichment (GSEA) scores we ranked the difference in (log) average expression of each gene between cases in the highest and lowest quartiles of *SRRT* expression for each cancer, using theses as ranked fold changes for input to GSEA using the *clusterProfiler* Bioconductor package^41^. GO terms were manually curated to remove uninformative redundancies.

### Statistical analysis

Statistical analysis were performed using GraphPad – Prism Version 9, or R v > 4.0. Statistical details can be found in the figure legends of corresponding experiments. For bioinformatic data processing, statistical analysis are detailed within the method description.

## Results

### Cytoplasmic isoforms of *Ars2*

The *Ars2* (*Srrt)* locus produces twelve transcripts in mouse and human, some of which encode different proteins isoforms (NCBI Gene database). In mouse, the transcripts can be organized into 3 groups based on potential protein products. The first group represents canonical ARS2, which initiates translation within exon 2 and encodes similar isoforms of 864-875 amino acids (Fig. 1A). This group of isoforms, which we designate ARS2n due to its nuclear localization, is well studied as a component of the CBC and has a role in RNA polymerase II (RNAP II) transcript biogenesis and turnover. The second group (2*) comprises 1 isoform, initiates translation within exon 4 and encodes a protein of 778 amino acids. The third group (3*) is comprised of 7 transcripts and involves alternative splicing of intron 5. Group 3* transcripts retain a portion of intron 5, are predicted to begin translation within either intron 5 or exon 6 and encode putative isoforms of 710 or 648 amino acids. Group 3* isoforms are especially intriguing since they are missing up to 227 amino acids from the functionally important N-terminus of ARS2n. This missing region contains a structural N-terminal helix-turn-helix, conserved tyrosine phosphorylation sites, and a nuclear localization signal. For simplicity, we designated the isoforms in group 3* as ARS2c (1, 2), for their potential to encode cytoplasmic isoforms of unknown function. Intron 5 is highly conserved across vertebrates and it is striking that 7 of the 12 *Ars2* isoforms retain portions of this intron (Fig. 1A).

**Fig. 1:**
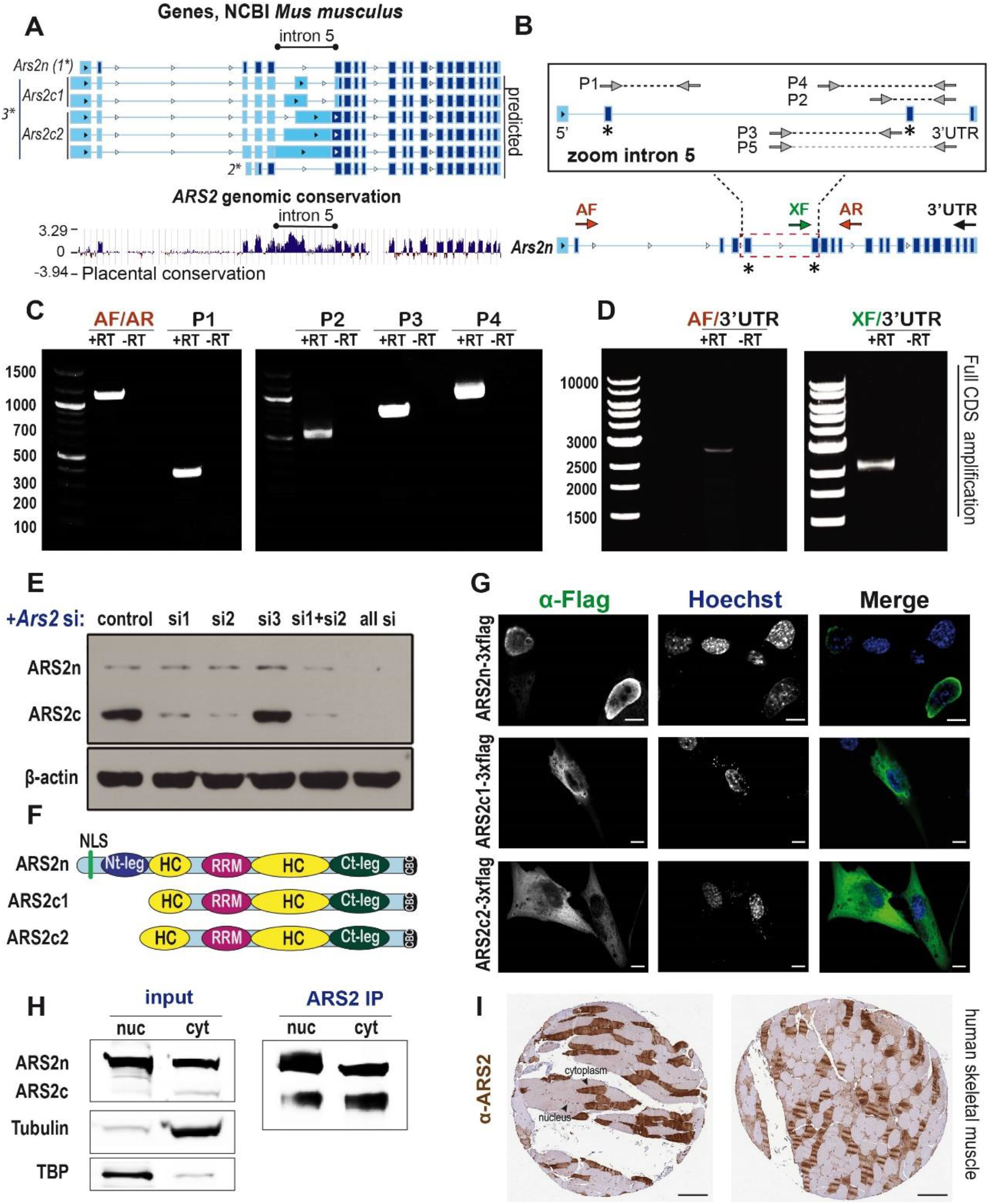
Cytoplasmic isoforms of *Ars2*. **A)** Representative *Ars2* sequences of major isoforms in NCBI Gene database (*mus musculus*), exons are represented in dark blue boxes. Bottom: conservation analysis of *Ars2* between mouse and 60 vertebrates. The alignments were generated using Multiz and UCSC/Penn State Bioinformatics comparative genomics alignment pipeline. Evolutionary conservation was measured using PhyloP and PhastCons. **B)** Diagram of primers used in **C)** and **D)** to amplify intron 5 **(C)** or the full CDS of the new isoforms **(D)**, in C_2_C_12_ cells. **E)** Western Blot of C_2_C_12_ whole cell lysates transfected with *Ars2* si or control. ARS2 was detected using an anti-ARS2 antibody and β-actin is shown as loading control. **F)** Protein motif structure of ARS2n and ARS2c1/c2 isoforms, based on the reported structure of ARS2n^20^. **G)** HEK 293T cells were transfected with ARS2n/c1/c2-3xflag. ARS2 was detected with anti-flag antibodies (green) and nuclei were stained with Hoechst 33342 (blue). Scale bar = 10 μM. **H)** Nuclear/cytoplasmic fractionation of C_2_C_12_ lysates. Endogenous ARS2 was immunoprecipitated and detected with anti-ARS2 antibodies. TBP and Tubulin detection are used as fractionation quality control. **I)** Immunohistochemistry image from Human Protein Atlas. Transverse cut of from skeletal muscle fibers. ARS2 was detected with anti-ARS2 antibody (HPA042858) shown in brown. Scale bar = 200μM. (https://www.proteinatlas.org/ENSG00000087087-SRRT/tissue/skeletal+muscle).

To investigate the expression of the group 3* transcripts, we first performed RT-PCR using combinations of primers flanking intron 5 (Fig. 1B). Indeed, we detected expression of transcripts retaining intron 5 in C_2_C_12_ myoblasts. Controls lacking reverse transcriptase did not show any amplification products confirming that our results are not a consequence of genomic contamination (Fig. 1C). Using a forward primer within intron 5 and reverse primer within the 3’UTR of *Ars2*, a full-length amplicon of 2500 bp was amplified (Fig. 1D). Sequencing showed that this transcript corresponded to variants XM_030254927.2, XM_006504631.3 and XM_036165632.1, which retain the second half of intron 5 and share the rest of the *Ars2* sequence from exon 6 until the 3’UTR. All amplicons detected in Fig. 1C and D were sequenced and were ≥99% identical to the sequences within the NCBI Gene Database (Suppl. sequence).

We next determined if the *Ars2c* isoforms are translated. Using an antibody (XL14.1) raised against the C-terminus of ARS2 (676-871), which is shared between ARS2n and ARS2c isoforms, we detected approximately 130 kDa and 100 kDa products in lysates from C_2_C_12_ cells. To confirm the identity of the products, we performed RNAi. Two of three unique siRNAs, targeting regions in common to all transcripts, knocked down expression of both protein products. The lower MW product showed increased sensitivity to the siRNA suggesting it is not a protein degradation product of the larger isoform. A combination of 3 siRNAs strongly knocked down both products (Fig. 1E). The results were confirmed using the Human Atlas antibody HPA042858, previously validated for detecting ARS2 by Western Blot. HPA042858, like XL14.1, detected 2 protein products (130-100KDa) that are knocked down when shRNA targeting all *Ars2* forms is used (Suppl. Fig. 1 A, B, C). These data strongly suggest that both ARS2n and ARS2c isoforms are translated in C_2_C_12_ cells.

To determine the localization of ARS2c isoforms, we expressed the cDNAs fused in-frame with a 3xFlag epitope. ARS2n isoforms have been shown to be predominantly nuclear but capable of shuttling between the nucleus and the cytoplasm^3^. As expected, ARS2n was nuclear in localization. In contrast, ARS2c1 and ARS2c2 localized to the cytoplasm (Fig. 1G). Similarly, nuclear/cytoplasmic fractionation and immunoprecipitation from C_2_C_12_ cell lysates showed that endogenous ARS2n and ARS2c are preferentially enriched in the nuclear and cytoplasm fractions, respectively (Fig. 1H).

The expression of cytoplasmic ARS2 isoforms is conserved in humans. In humans, an isoform analogous to ARS2c is encoded by XM-024446794.1 (SRRT-X8) (Suppl. Fig. 2A). Using RT-PCR, Western Blotting, and shRNAs specific for all *ARS2* isoforms or intron 5 exclusively, we demonstrated that ARS2c is expressed in the human cell line HeLa (Suppl. Fig. 2B, C, D, E). Additionally, the Human Protein Atlas shows that ARS2 localizes to both the nucleus and cytoplasm in several tissues including muscle, implying expression of both ARS2n and ARS2c isoforms (Fig. 1I). While we cannot rule out some of the cytoplasmic immunofluorescence signal is due to shuttling of the ARS2n, taken together, our data strongly indicates the presence of cytoplasmic isoforms of ARS2.

### Comparison of ARS2n and ARS2c interactomes

To understand the biological roles of ARS2c in comparison to nuclear ARS2n, we analyzed their interactomes using BioID and LC-MS/MS. Proximity-dependent biotin identification (BioID) allows the detection of transient and weak protein interactions in living cells^35,42^. In BioID, a promiscuous bacterial biotin ligase (BioID2) is fused to the protein of interest. After the addition of the biotin substrate, proximal proteins are biotinylated and subsequently purified by streptavidin capture (Fig. 2A). Biotinylated proteomes were normalized against the ligase control and enriched proteins were detected by Western Blot or LC-MS/MS (Fig. 2B). We confirmed that fusion of ARS2 isoforms to BioID2 ligase did not affect their localization or functionality: BioID2-ARS2n and BioID2-ARS2c1/c2 localized in the nucleus and cytoplasm respectively, and BioID2-ARS2n maintains its interaction with the CBC complex (Suppl. Fig. 3A, B). Interactome enrichment was higher when the biotin ligase enzyme was placed on the N-terminus of ARS2 versus the C-terminus, which likely reflects the importance of the CBC binding site within the extreme C-terminus of ARS2 (data not shown). We generated Flp-In T-REx 293 cell lines, in which the expression of ARS2 isoforms or biotin ligase control could be induced and regulated with the addition of tetracycline. However, this method returned very few enriched interactors, because we could not adequately control the levels of the biotin ligase control relative to ARS2c1,c2 expression. However, we did confirm ARS2n interactions with known interactors: NCBP3, ZC3H18, ZC3H4, THOC2 and PHAX using this method. To increase the enrichment of ARS2 isoforms interactomes to levels comparable to the literature^20^ we overexpressed ARS2 isoforms and control. An equal amount of protein for each condition, as quantified by silver stain, was used for LC-MS/MS.

**Fig. 2:**
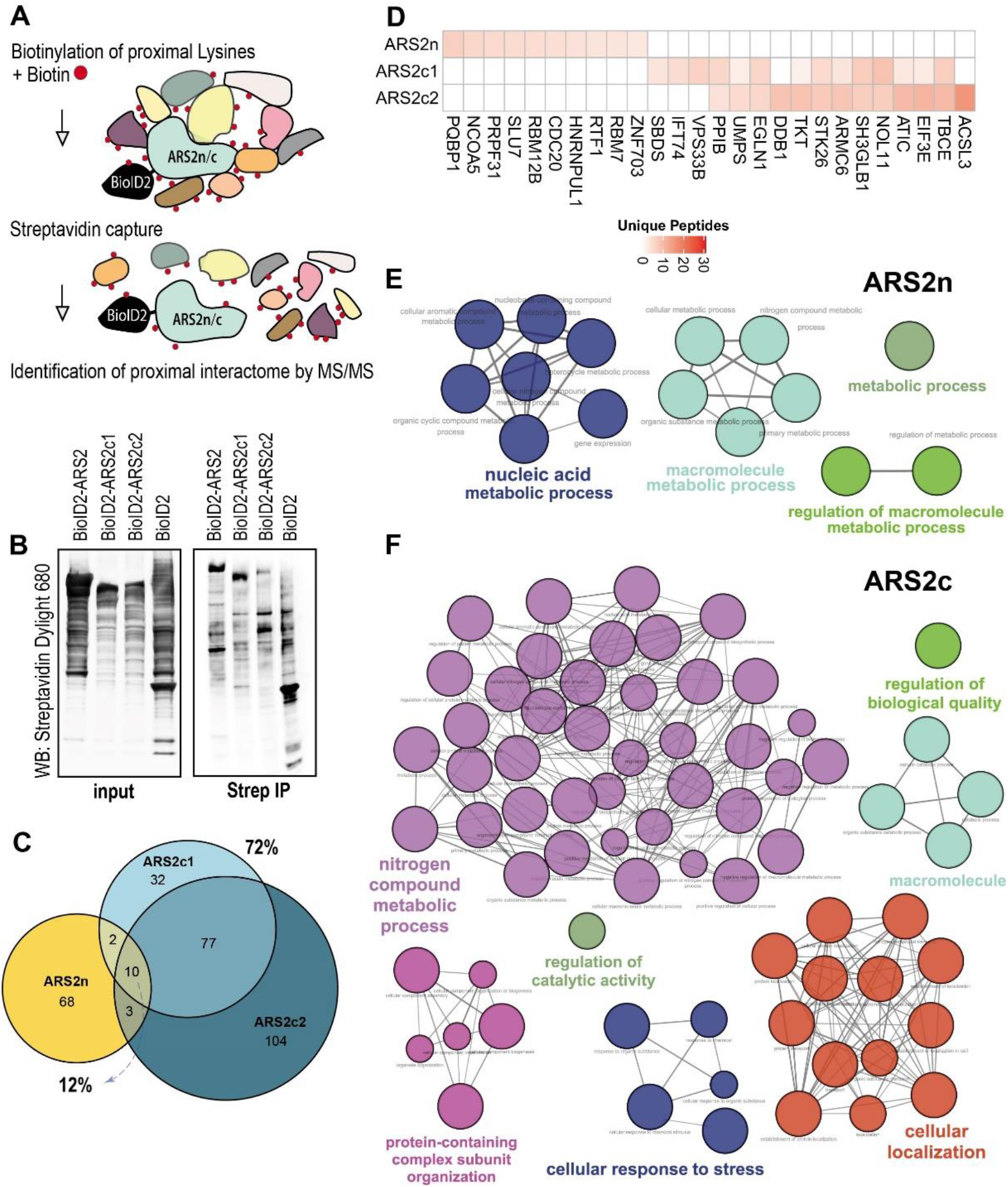
Comparison of ARS2n and ARS2c interactomes. **A)** Schematical representation of BioID. Biotynilated proteins are immunoprecipitated by streptavidin capture and interactors are detected by Western Blot or LC/MS-MS. **B)** HEK 293T cells were transfected with BioID2-ARS2n/c1/c2-3xflag or control, and treated with biotin for 24h. After lysis, biotinylated interactomes of ARS2 forms and BioID2 control are immunoprecipitated with streptavidin beads and detected by western blot using a streptavidin dylight 680 antibody. **C) D) E) and F)** Samples were treated as in **B)**, protein interactomes were detected by LC/MS-MS. For each condition,15 biological replicates were pooled and ran in 3 independent experiments. ARS2 interactomes were normalized against BioID2 control and analyzed by SAINT. Only proteins with a SAINT score (SP>0.7) were included in the subsequent analysis. **C)** Venn diagram comparing ARS2 forms. **D)** Top ten interactors for each isoform. **E) F)** Gene ontology (GO) analysis of ARS2 isoform interactomes using ClueGO^37^. The size of the nodes reflects term significance after Bonferroni correction. Only GO terms with a p.value < 0.01 were considered significant and therefore represented in the network.

The proximity interactome of ARS2n confirmed known interactors including NELF, NCBP3, PHAX, XRN2 and RBM7, previously detected by AP-LC-MS/MS^19,20^ (Suppl. Fig. 3C). As expected, given the difference in localization, ARS2n shared only 12% of its interactome with ARS2c isoforms. In contrast, 72% of the ARS2c1 interactome is shared by ARS2c2 (Fig. 2C). Indeed ARS2c isoforms share most of their top 10 interactors (Fig. 2D). Due to the similarities between ARS2c isoforms interactomes we decided to focus on ARS2c2, as it had a larger number of positive interactors, and refer to it as ARS2c, for simplicity. Global network analysis of the ARS2n interactome shows an enrichment for proteins related to nucleic acid and macromolecule metabolism, which is consistent with the role of ARS2n in RNA metabolism (Fig. 2E). The ARS2c interactome, on the other hand, is enriched in proteins that participate in the biological processes of nitrogen compound metabolic processes, cellular localization, and cellular response to stress (Fig. 2F). This last group is intriguing since the *Ars2* gene has been reported to confer sensitivity to the cellular stressor arsenic^3^.

### ARS2c and the cellular response to arsenic stress

Arsenic treatment suppresses global transcription and translation, and in our case caused a notable shift in the proteome of cells and a dramatic reduction of protein diversity. After arsenic treatment, there is an accumulation of proteins between 50-75KDa (Fig. 3A). Consistent with ARS2c being important for arsenic sensitivity, expression of ARS2c is upregulated in response to arsenic, whereas expression of ARS2n, and known interactors of ARS2: NCBP1 and UPF1, are downregulated (Fig. 3B, C). The induction of ARS2c expression appears to be at the transcription or RNA stability level and was inhibited by siRNA against ARS2c (Fig. 3D). Full sequences of RNAi and shRNAs used in the study, validation, and targeting diagrams are included in Suppl. table 1, Suppl. Fig. 4 A-C and 2C. We tested other cellular stressors such as tunicamycin, serum starvation, and translation inhibition (Puromycin) (Suppl. Fig. 5 A-D). Interestingly, only puromycin treatment induced ARS2c expression (Suppl. Fig. 5 C-E).

**Fig. 3:**
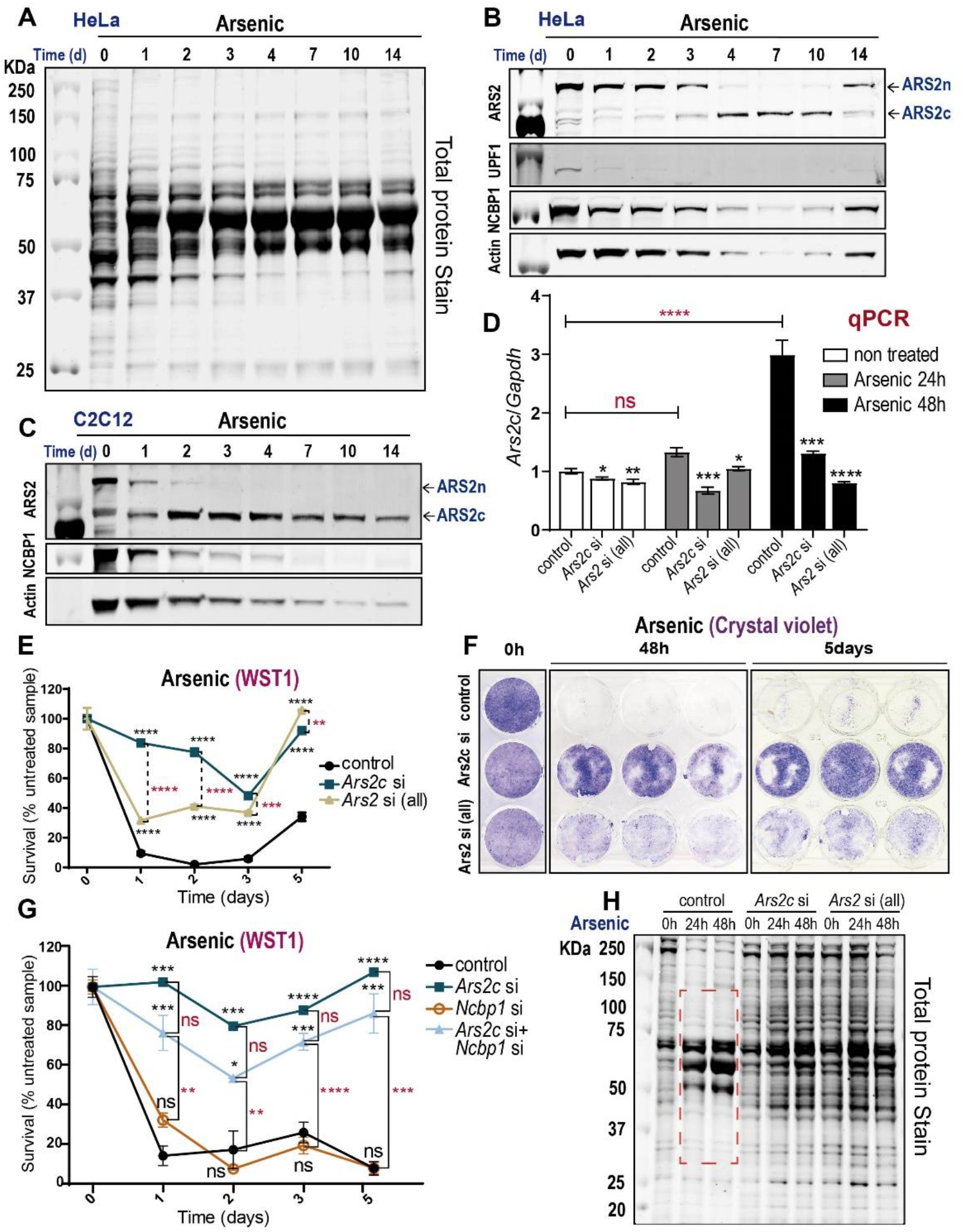
ARS2c required for arsenic stress response. **A) B) C)** HeLa and C_2_C_12_ cells were treated with 40uM arsenic trioxide for 14 days. Western blots of the cell lysates are shown with the indicated antibodies. Total protein was detected by Revert™ 700 Total Protein Stain. **D)** C_2_C_12_ cells transfected with *Ars2c*/*Ars2*(all)/control RNAi were treated with arsenic trioxide for 48h. *Ars2c* was detected by RT-qPCR, using a primer against intron 5. Data are represented as mean ± SEM for n=3 biologically independent samples. Statistical analysis: One way ANOVA, post-test Tukey’s (*p≤0.05; **p≤0.01; ***p≤0.001, ****p≤0.0001). **E) F) G)** C_2_C_12_ cells transfected with *Ars2c*/*Ars2*(all)/*Ncpb1*/*Ars2c*+*Ncbp1*/control RNAi were treated with arsenic trioxide for 1-5 days. Survival (% treated/untreated sample) was measured by WST1 **(E, G)** or crystal violet **(F)**. Data are represented as mean ± SEM **(E, G)** for n=3 biologically independent samples for each time point. Statistical analysis: One way ANOVA, post-test Tukey’s (*p≤0.05; **p≤0.01; ***p≤0.001, ****p≤0.0001). **H)** C_2_C_12_ cells were transfected with *Ars2*c/*Ars2*(all)/control RNAi and treated with arsenic trioxide for 48h. Cell lysates were analyzed by Western blot and total protein was detected as in **(A)**. Each lane represents 3 pooled biologically independent samples.

To evaluate the importance of ARS2c upregulation during arsenic treatment, we knocked down ARS2c and examined cellular survival in response to arsenic, by measuring metabolically active cells using the WST-1 assay. Depletion of cytoplasmic ARS2 (*Ars2*c si, blue) significantly increases the resistance of C_2_C_12_ myoblast cells to arsenic treatment (Fig. 3E). Depletion of all ARS2 isoforms (*Ars2*all si, gray), also increases the resistance of C_2_C_12_ to arsenic but to a lesser extent than ARS2c knockdown alone. Since ARS2c is similarly knocked down by *Ars2* si (all) and *Ars2*c si (Fig. 3D), the significantly different phenotype of cells exposed to these RNAi suggests that ARS2n may be playing an opposite role in the pathway, providing resistance to arsenic. We confirmed the effect of ARS2c on arsenic sensitivity using a crystal violet assay (Fig. 3F, Suppl. Fig. 5F).

Surprisingly, the ARS2c mediated arsenic response seems to be independent of the CBC complex. Downregulation of NCBP1 does not increase cellular survival following arsenic treatment, and co-depletion of ARS2c and NCBP1 results in an arsenic resistant phenotype comparable to depletion of ARS2c alone (Fig. 3G, Suppl. Fig. 5H, I). Additionally, *Ars2*c knockdown had no effect on the survival following puromycin treatment (Suppl. Fig. 5G), indicating that the effect of ARS2c on arsenic sensitivity is not due to the ability of both arsenic and puromycin to block translation.

The wholescale changes in the proteome, which occur in response to arsenic, suggest that this change is important for the cellular response to the chemical. We asked whether ARS2c is required for proteomic remodelling in response to arsenic. Consistent with the importance of ARS2c in the arsenic sensitivity, downregulation of ARS2c attenuates the proteomic remodelling (Fig. 3H). Previous studies implicating the role of *Ars2* in arsenic sensitivity examined the effects of *Ars2* knockdown on sensitivity^3^. Since the isoforms were not known, it was assumed the observed effects were due to ARS2n. Our data now show that it is the ARS2c isoforms and not ARS2n that confer arsenic sensitivity on cells. The effects of ARS2c on arsenic sensitivity are independent of the CBC complex and the ability of arsenic to inhibit translation. Instead, ARS2c is upregulated in response to arsenic and is necessary for the cellular response and sensitivity to arsenic stress.

### The role of ARS2n and ARS2c in NMD

ARS2n is a scaffold protein that joins the CBC to direct RNAPII transcript processing, degradation, and export. To better understand the role of ARS2c isoforms, we asked whether they share functions with ARS2n. GO analysis shows that ARS2n and ARS2c isoforms share interactions with components of the mRNA-RNA catabolic process: NMD (Fig. 4A). Consistent with previous reports, we detected proximity interactions of ARS2n with core EJC components MAGOH, MAGOHB and CASC3 (Fig. 4B, left). In contrast, ARS2c isoforms are enriched for NMD components UPF1 and ERF1 (Fig. 4B, right). We confirmed the novel interactions between ARS2c and ERF1, UPF1, as well as components of the translation initiation complex EIF3K/E/F, by immunoprecipitation and western blotting (Fig. 4C,D). We know from previous work that ARS2n interacts with all four core components of the EJC: CASC3, MAGOH, RBM8A and EIF4A3, the EJC periferal factors: UPF3B, RNPS1, PNN and ACINUS as well as UPF1 and the phosphatases PPP2R1A and PPP2R2A (Fig. 4E)^5,19,20,43–46^. Additionally, the CBC complex (NCBP1/2) has been shown to stabilize the interaction between UPF1-ERF1/ERF3 and the EJC, promoting NMD^9,18^. However, to our knowledge no role for ARS2 in NMD has been described and this is the first report of specific interactors for ARS2c isoforms with this pathway.

**Fig. 4:**
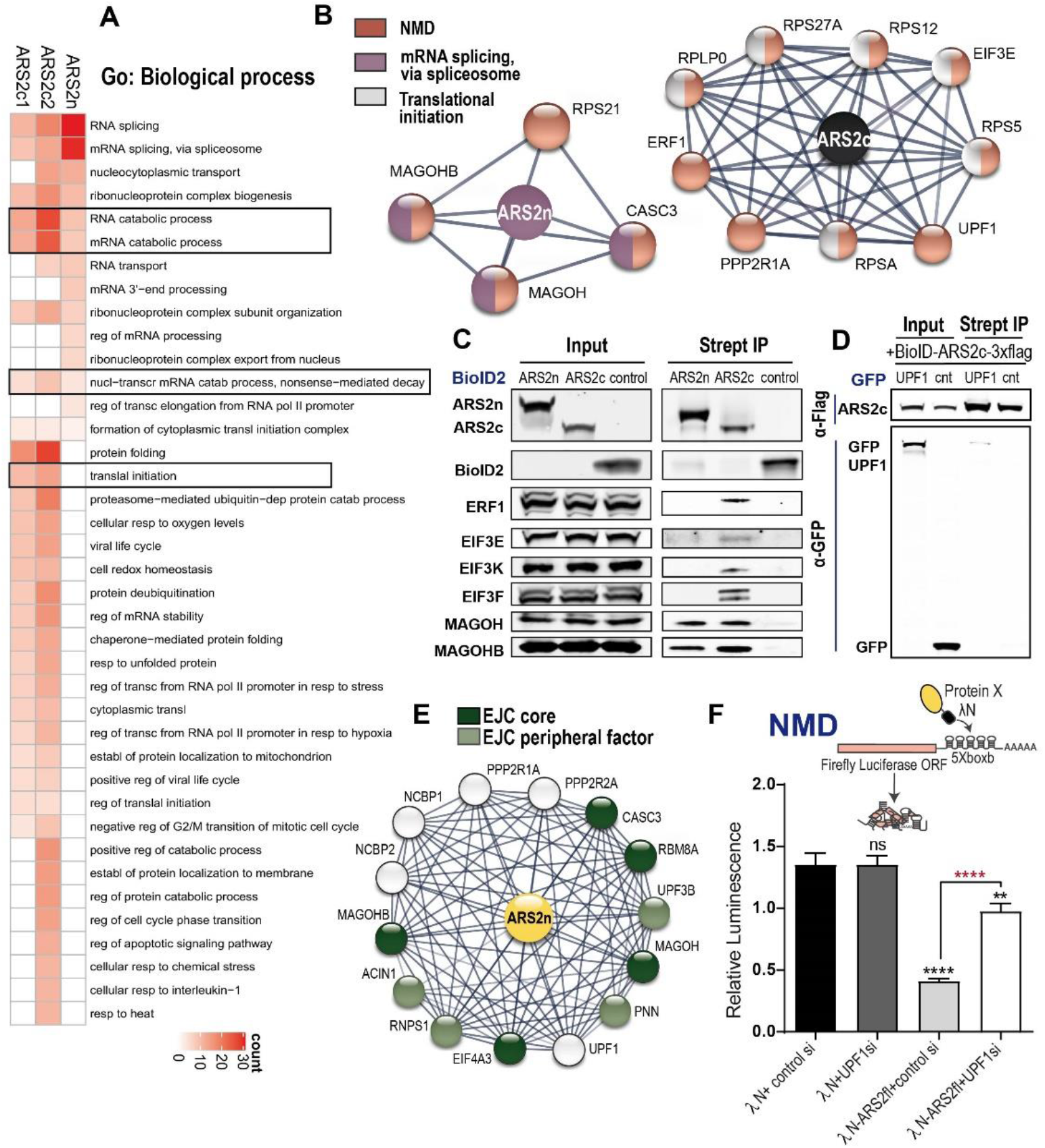
ARS2 isoforms interact with different components of NMD pathway. **A)** Gene Ontology (GO) analysis of ARS2 isoform interactomes. Enriched biological process are shown (p.adjust < 0.05). **B)** NMD pathway components interacting with the isoforms are represented using STRING^59^. **C) D)** Western Blot confirmation of detected interactors using the specified antibodies. **C)** HEK 293T cells were transfected with BioID2-ARS2n/c1/c2-3xflag or control and treated with biotin for 24h. Biotinylated interactors were pulled down with streptavidin beads and proteins were detected as indicated. **D)** HEK 293T cells were co-transfected with BIOID2-ARS2c-3xflag and GFP-UPF1. Samples were pulled down using anti-Flag magnetic beads and UPF1 was detected using an anti-GFP antibody. **E)** STRING representation of the interactions between ARS2n and NMD components, reported in the literature. **F) Top:** Schematic representation of the tethering assay. **Bottom:** HeLa cells were transfected with λN/λN-ARS2n + control si/*UPF1* si + Firefly luciferase reporter + Renilla luciferase. Normalized Luciferase activity was measured at 24h after transfection. Data are represented as mean ± SEM for n=9 biologically independent samples. Statistical analysis: One way ANOVA, post-test Tukey’s (**p≤0.01; ****p≤0.0001).

To evaluate the role of ARS2n in NMD, we initially performed a tethering assay in which the protein of interest is fused with a λN tag and binds to a 5x-B-Box in the 3’ UTR of a luciferase reporter (Fig. 4F top, Suppl. Fig. 6A). The tethering assay mimics the deposit of an EJC downstream of a stop codon and measures the ability of a protein of interest to recruit components of the NMD pathway and trigger decay of the reporter mRNA. Tethering of ARS2n to the luciferase reporter resulted in reporter degradation (Suppl. Fig. 6B). To test whether the observed degradation was dependent on NMD, we knocked down *UPF1*, a key component of the NMD pathway. As shown in Fig. 4F, knockdown of *UPF1* partially rescued the expression of the reporter, indicating the effect of ARS2n is dependent on UPF1. Together with our BioID analysis, these data suggest a role for ARS2 isoforms in NMD.

To gain a better understanding of the role of the nuclear and cytoplasmic ARS2 isoforms in NMD, we used a NMD fluorescent reporter^28^. The pNMD+ vector contains GFP cDNA fused to exon 2 and exon 3 of human β-globin. Splicing of the β-globin intron deposits an exon junction complex (EJC) >50 nucleotides downstream of the termination codon of GFP. Consequently, the GFP termination codon is recognized as a premature termination codon (PTC) and GFP mRNA is degraded by NMD. The pNMD-vector is a negative control. In this plasmid, exon 2 is only 35 nucleotides long. After splicing, the distance between the termination codon of GFP and the EJC will be <50 nucleotides and therefore insensitive to NMD. To normalize for transfection efficiency, the vectors also contain an expression cassette for the far-red protein Katushka (TurboFP635) (Fig. 5A, B, Suppl. Fig. 6C). For each transfection, GFP is normalized to Katushka and then expressed as the ratio of pNMD-ve/pNMD+ve. Therefore, GFP stabilization during impaired NMD results in a smaller pNMD-ve/pNMD+ve ratio.

**Fig. 5:**
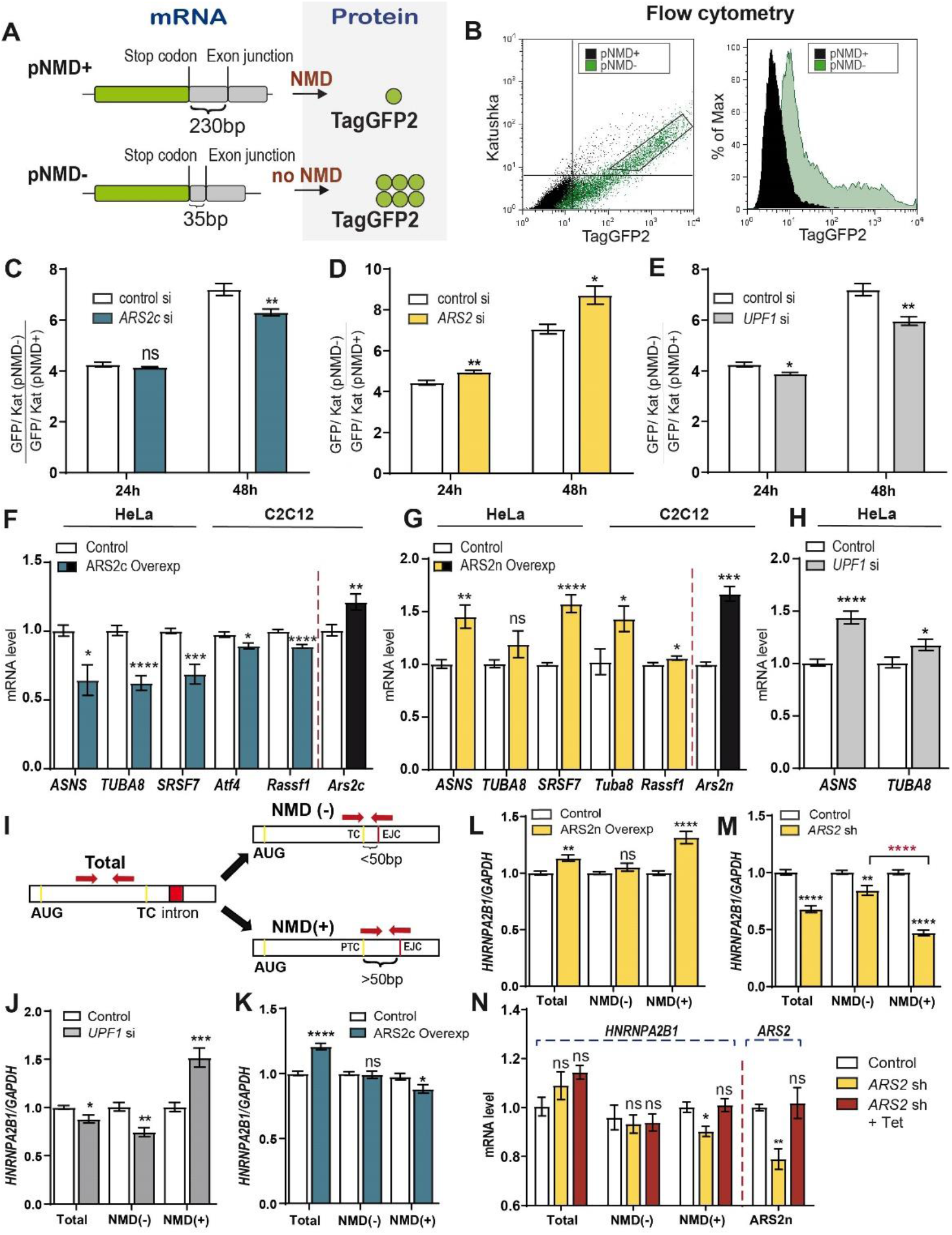
The role of ARS2n and ARS2c in NMD. **A)** Schematic representation of pNMD+/pNMD-reporters. pNMD+ is degraded by NMD, while pNMD- is insensitive to NMD degradation. **B)** Flow cytometry showing pNMD+ degradation and reporter functionality. **C) D) E)** HeLa cells were transfected with pNMD+/pNMD- and *ARS2c*/*ARS2*/*UPF1* or control RNAi. Expression levels were detected by Flow cytometry. TagGFP2 levels were normalized to Katushka, shown are the pNMD-/pNMD+ ratios. Data are represented as mean ± SEM for n=3 or n=6 biologically independent samples. Statistical analysis: two-tail unpaired t-test (*p≤0.05; **p≤0.01). **F) G) H) J) K) L) M)** RT-qPCR performed after transfection of HeLa and C_2_C_12_ cells with ARS2c-GFP/ARS2n-GFP*/*GFP or *ARS2* sh/*UPF1*si/negative controls for 48h. Each gene was normalized against *GAPDH*. **I)** Schematic representation of primers used **J) K) L) M)** and **N). N**) RT-qPCR of Flp-In T-REx 293 cells inducibly expressing *ARS2*n were transfected with *ARS2* sh or control for 48h and treated with 2.27ng/ml of tetracycline for 36h to induce *ARS2n* to endogenous levels. Data are represented as mean ± SEM for n=3 to n=6 biologically independent samples (see Materials and Methods section). Statistical analysis **F) G) H) J) K) L) M)** two-tail unpaired t-test (*p≤0.05; **p≤0.01; ***p≤0.001, ****p≤0.0001), **N)** One way ANOVA, post-test Dunnett’s (*p≤0.05; **p≤0.01).

Using this system, we found that ARS2c and ARS2n have opposing impacts on NMD. Specific knockdown of *ARS2*c, promotes the accumulation of the pNMD+ reporter with respect to the pNMD-control, as measured by flow cytometry quantification of GFP protein and RT-qPCR quantification of the GFP transcript at 48h (Fig. 5C, Suppl. Fig. 6F). A similar effect was observed with downregulation of UPF1 (Fig. 5E, Suppl. Fig. 6H), suggesting that ARS2c isoforms are required for NMD. We confirmed the role of ARS2c in NMD is conserved in mouse cells by testing this reporter assay in C_2_C_12_ cells. A similar stabilization of the pNMD+ GFP protein and pNMD-/pNMD+ ratio decrease was seen in these cells (Suppl. Fig. 6D). There are no sequences unique to *ARS2*n to specifically target this isoform. Thus, to test the impact of ARS2n on NMD we used RNAi to exon 4, which disrupts expression of both *ARS2*n and *ARS2*c (Suppl. Fig. 2C). Surprisingly, this RNAi had the opposite effect, destabilizing the pNMD+ reporter relative to the pNMD-control, at both protein and transcript levels (Fig. 5D, Suppl. Fig. 6G). Collectively, these experiments suggest that ARS2n inhibits the NMD pathway, while ARS2c promotes it.

To confirm the opposing roles of ARS2n and ARS2c in NMD, we next examined the effect of isoform overexpression and downregulation on endogenous genes that are naturally regulated by NMD. Consistent with a promoting role in NMD, ARS2c overexpression increased the activity of the NMD pathway and promoted degradation of the endogenous NMD targets (Fig. 5F). Conversely, both UPF1 and ARS2c depletion inhibited the activity of the NMD pathway and favored the accumulation of endogenous NMD targets (Fig. 5H, Suppl. Fig. 6I). These results confirm that ARS2c, similar to UPF1, promotes the NMD pathway. In contrast, overexpression of ARS2n decreased the activity of the NMD pathway and induced the accumulation of NMD regulated transcripts (Fig. 5G). Consistent with an inhibitory role, downregulation of ARS2n increased NMD activity and promoted the degradation of endogenous NMD targets (Suppl. Fig. 6J). These results were observed in both human HeLa and mouse C_2_C_12_ cells, confirming the effects of the isoforms on endogenous targets are conserved in between the species.

We next demonstrated that the effect of ARS2 isoforms on the pathway was NMD-dependent and not due to off-targeting. We used a set of primers to amplify alternatively spliced transcripts of the *HNRNPA2AB1* gene^47^ in HeLa or in a Flp-In T-REx 293 inducible cell line, in which 3xFlag-ARS2n has been knocked into an *FRT* locus within the cell line. The endogenous *HNRNPA2AB1* gene generates two alternative splicing transcripts represented here as NMD- and NMD+ (Fig. 5I). Similar to the pNMD-/+ reporter, the NMD-transcript is insensitive to NMD, while the NMD+ product is degraded by NMD. The total transcript levels are measured by an amplicon common to all the splicing variants. As expected, downregulation of UPF1 results in an accumulation of the NMD+ transcript (Fig. 5J). As shown above, overexpression of ARS2c preferentially induces the degradation of the NMD+ transcript (Fig. 5K). In contrast, overexpression of ARS2n promotes the accumulation of the NMD+ transcript, while downregulation of ARS2n preferentially induces NMD+ degradation (Fig. 5L, M). To confirm that the effects of *ARS2* knockdown on the pathway were specific to *ARS2n*, we repeated the experiment but induced expression of 3xFlag-ARS2n with the addition of tetracycline. Restoring *ARS2*n levels rescued the expression of the HNRNPA2AB1 NMD+ transcript, showing that the inhibitory effect of *ARS2n* is specific. (Fig. 5N).

### ARS2 isoforms work in tandem to regulate NMD

To understand how ARS2 isoforms differentially regulate NMD, we first asked whether the inhibitor role of ARS2n on the NMD pathway was dependent on its nuclear localization. We narrowed the nuclear localization signal region from 105 amino acids^48^ to 12, and generated an ARS2n mutant which lacks amino acids (73-LSPPQKRMRRDW-84). As shown in Fig. 6A, deletion of these 12 amino acids localizes ARS2n to the cytoplasm. Interestingly, deletion of the nuclear localization signal (NLS) was sufficient to abrogate the inhibition of NMD caused by ARS2n overexpression, suggesting that the inhibitory effects of ARS2n on the pathway originate in the nucleus (Fig. 6B).

**Fig. 6:**
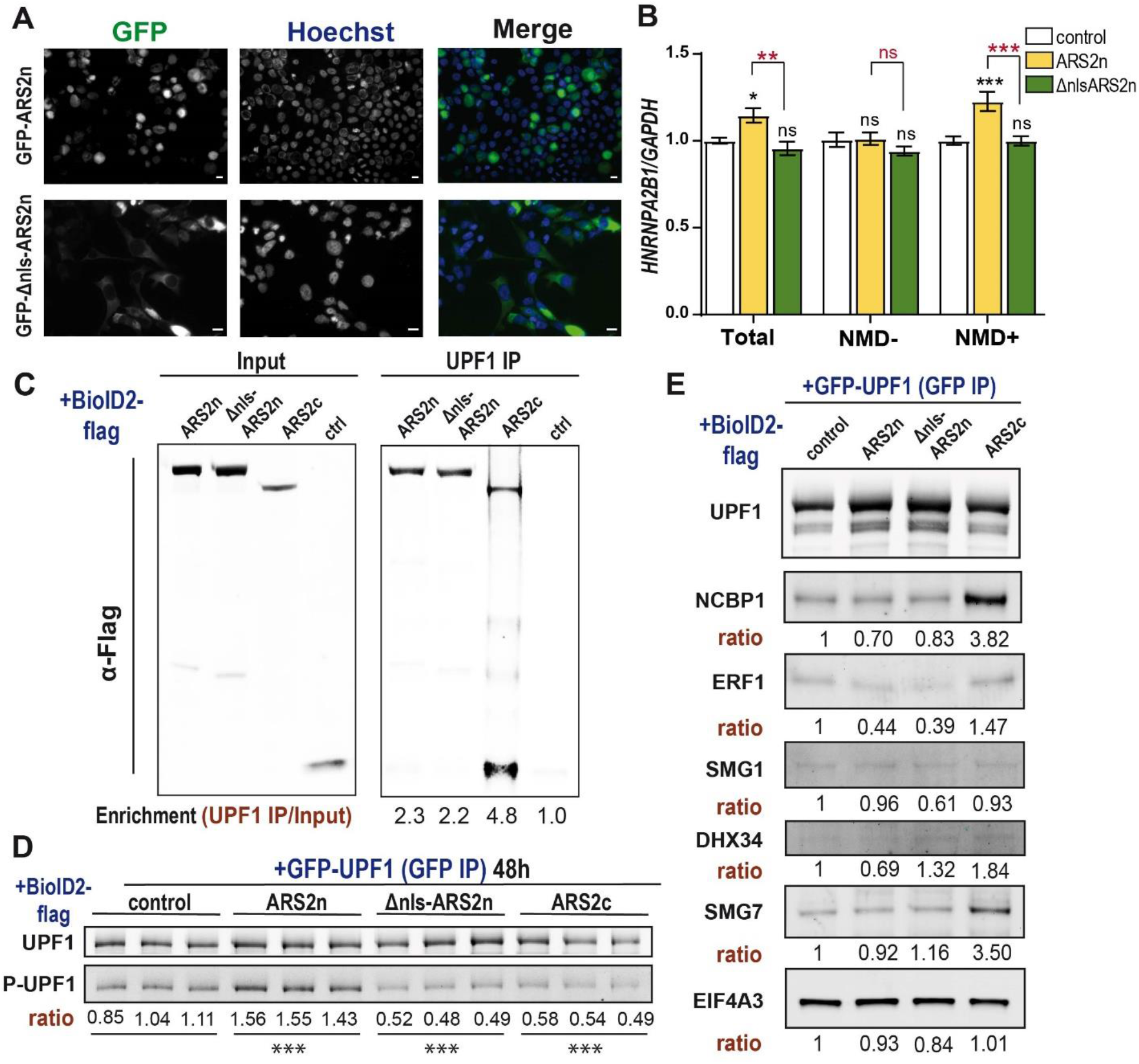
ARS2 isoforms work in tandem to regulate NMD. **A)** HEK 293T cells were transfected with GFP-ARS2n/ GFP-Δnls-ARS2n. GFP fusion proteins are shown in green and nuclei were stained with Hoechst 33342 (blue). Scale bar = 10 μM. **B)** RT-qPCR performed after transfection of HeLa cells with GFP-ARS2n/ GFP-Δnls-ARS2n or GFP control for 48h. *HNRNPA2AB1* gene isoforms were detected using Fig. 5I primers. Each gene was normalized against *GAPDH*. Data are represented as mean ± SEM for n=6 biologically independent samples. Statistical analysis: One way ANOVA, post-test Tukey’s (*p≤0.05; **p≤0.01; ***p≤0.001). **C)** HEK 293T cells were transfected with BIOD2-ARS2n/Δnls-ARS2n/ARS2c/control-3xflag. Endogenous UPF1 was pulled down with anti-UPF1 antibody and ARS2n/Δnls-ARS2n/ARS2c/control were detected using anti-flag antibodies. Enrichment is defined as the ratio between ARS2n/Δnls-ARS2n/ARS2c/control in the pull-down, versus input. **D) E)** HEK 293T cells were transfected with BIOD2-ARS2n/Δnls-ARS2n/ARS2c/control-3xflag and GFP-UPF1. Specified proteins are detected in GFP-UPF1 pull-downs. Ratios represent normalization to GFP-UPF1 in the sample. **D)** Three biologically independent samples are shown. Statistical analysis: One way ANOVA, post-test Dunnett’s (***p≤0.001).

To evaluate the mechanism of the isoforms and mutant Δnls-ARS2n on NMD, we next looked at their ability to interact with UPF1. All three ARS2 proteins interacted with UPF1. However, ARS2c was more enriched in UPF1 pull-downs than ARS2n or the ΔNLS mutant (Fig. 6C). Overexpression of ARS2 isoforms also differentially affected phosphorylation of UPF1. Overexpression of ARS2n increased UPF1 phosphorylation, while ARS2c and the ΔNLS mutant decreased UPF1 phosphorylation (Fig. 6D). We next evaluated the effect of the differential UPF1 interaction and phosphorylation on the activity of the NMD pathway. Consistent with an inhibitory role in NMD, overexpression of ARS2n decreased the interaction between UPF1 and NCBP1, ERF1, SMG1, DHX34 and SMG7 (Fig. 6E). In contrast, overexpression of ARS2c, increased the binding of UPF1 to NCBP1, ERF1, DHX34 and SMG7 (Fig. 6E). Interestingly the ΔNLS mutant did not promote the SURF complex formation as effectively as ARS2c (Fig. 6E). This result indicates that the ΔNLS mutant is not able to replace ARS2c, implying that the loss of the N-terminal leg of the protein is functionally important for NMD promotion.

### The NMD pathway is associated with ARS2 *in vivo*

Missregulation of ARS2 has pathological consequences. Analysis of gene expression and survival data for solid cancers publicly available through TCGA, revealed that total *ARS2 (SRRT)* expression varies both between tumour types and within the same tumour types (Suppl. Fig. 7). Expression of ARS2 is significantly associated with poorer survival in several cancers, including liver hepatocellular carcinoma, glioma, and kidney renal cell cancer (i.e. LIHC, LGG and KIRC; Cox proportional hazards models, P_adj_ < 0.05). In contrast, ARS2 expression predicted better outcome in pancreatic adenocarcinoma (PAAD; Fig. 7A, B). Since missregulation of NMD is also associated with tumour progression^49^, we decided to investigate the relationship between ARS2 and NMD in the tumour scenario.

**Fig. 7.**
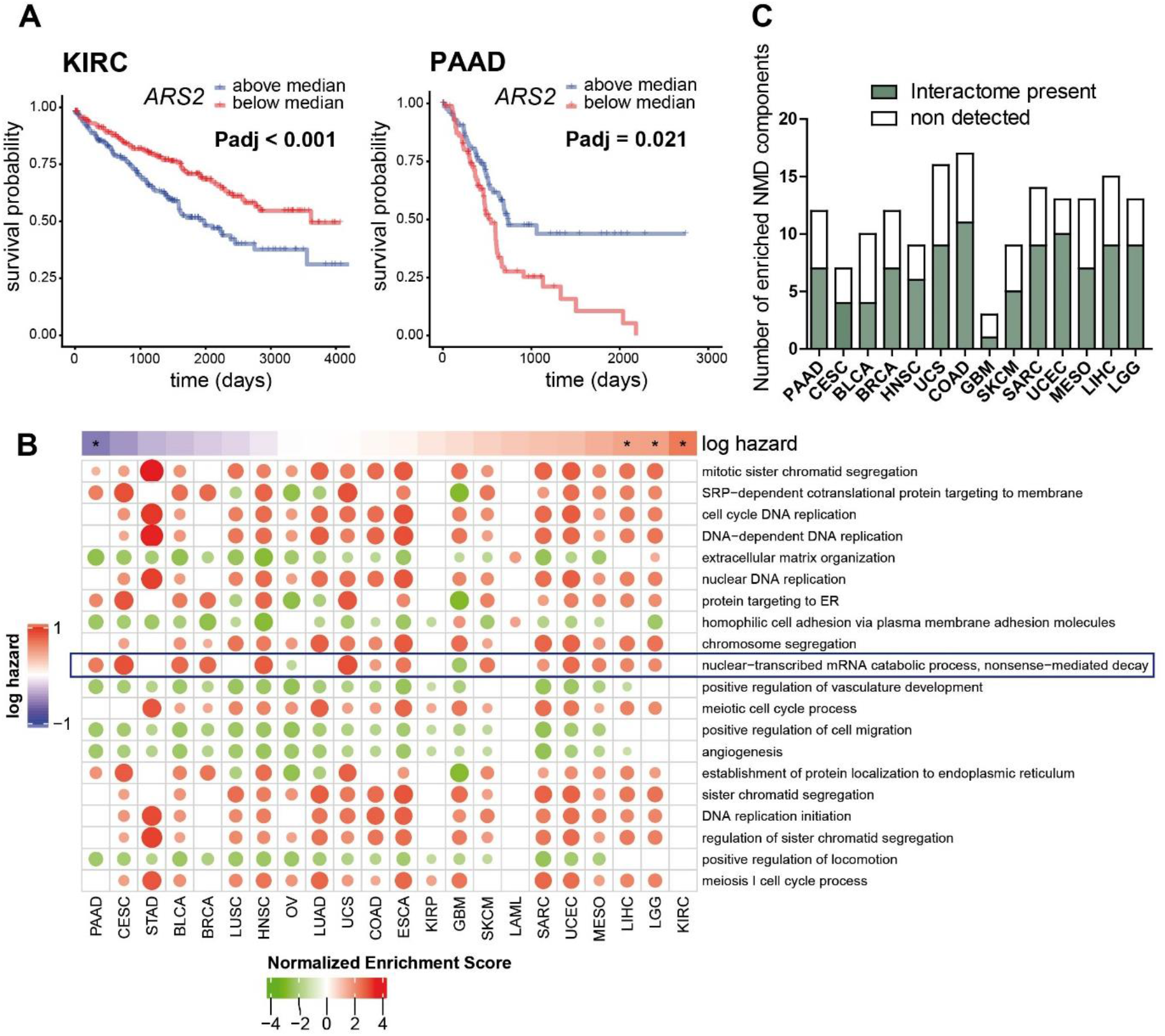
NMD pathway is enriched in high ARS2-expressing tumours. **A)** Kaplan-Meier survival plots for kidney renal clear carcinoma (KIRC) and pancreatic adenocarcinoma (PAAD) patients with high or low ARS2 expression. Data was obtained from the TCGA database. **B)** ARS2 survival data for solid cancers evaluated across tumours in the TCGA database. (Cox proportional hazards models, Padj<0.05 represented by asterisk). GSEA was performed using genes ranked-fold change of expression difference between patients with high (first quartile) and low ARS2 expression (fourth quartile) for each represented tumour. Top 20 GO terms associated with ARS2 expression (biological processes; based on average absolute normalized enrichment score (NES) across cancers) are shown; circle sizes are proportional to NES. **C)** NMD components enriched in the GSEA were compared against the ARS2n interactome reported in the literature. The ratio of enriched NMD components that are reported ARS2n interactors against total enriched NMD components is represented in green.

To evaluate which biological processes were correlated with expression of ARS2, we compared cancers in the first and fourth quartiles of *ARS2* expression for each evaluable cancer type and performed gene set enrichment analysis based on the ranked log-fold changes of gene expression between quartiles. Strikingly, this unbiased approach revealed that the NMD pathway is highly enriched in most of the evaluated tumour types. The top 20 GO terms associated with *ARS2* expression (biological processes; based on the average absolute normalized enrichment score across cancers) are shown in Fig. 7B. Moreover, analysis of the specific NMD components that contribute to this score across tumour types (Fig. 7C), shows that approximately half are confirmed interactors of ARS2 (Suppl. Table 6). The fact that the NMD pathway is enriched when ARS2 expression is elevated in diverse cancer types suggests a role for ARS2 isoforms in the regulation of the NMD pathway *in vivo* and raises important questions about the clinical relevance of the relationship between ARS2 and the NMD pathway in cancer.

## Discussion

In the current study, we have identified dedicated cytoplasmic isoforms of *Ars2* in human and mouse cell lines that function in the regulation of NMD, and in the cellular response to arsenic induced stress. We propose a model in which ARS2n and ARS2c work in tandem to regulate NMD (Fig. 8). As previously described, ARS2n primarily functions in the nucleus in conjunction with the CBC to promote mRNA splicing, degradation and export^1–7^. We propose that once in the cytoplasm, an isoform switch occurs in which ARS2c replaces ARS2n within the CBC. This ARS2 isoform “switching” has dramatic functional consequences: changing ARS2 from a NMD inhibitor to a NMD promoter that enhances SURF complex formation and transcript degradation. This model is consistent with our data showing that association of ARS2c with the CBC promotes the interaction of UPF1 with NCBP1, ERF1 and DHX34, the formation of the SURF complex at the PTC, SMG7 recruitment and transcript degradation though NMD. The model is also consistent with the known role of CBC-ARS2n as a platform for CBC dependent RNA processing in nucleus and with previous literature showing that NCBP1 promotes both UPF1 interaction with ERF1 and ERF3 and the association of UPF1 with the EJC^9,18^. We suggest that failure to switch ARS2 isoforms in the cytoplasm creates a CBC that is inefficient at promoting NMD. Thus, in the case of NMD, ARS2 isoform switching tailors the proteins to function in a nonredundant manner in both nuclear and cytoplasmic CBC-dependent processes.

**Fig. 8:**
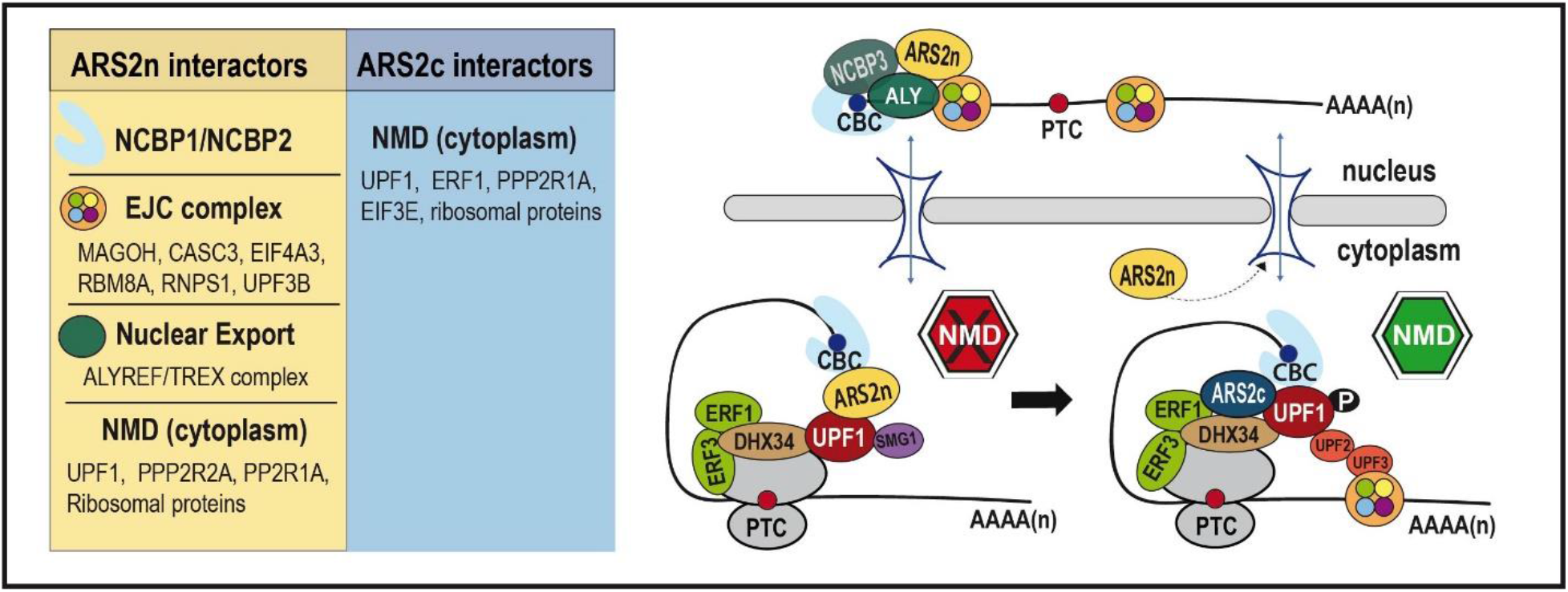
Model of ARS2n and ARS2c role in NMD. ARS2n, bound to NCBP1/2/3, interacts with the EJC and regulates the translocation to the cytoplasm of NMD-sensitive mRNAs. Once in the cytoplasm, ARS2n is recycled back to the nucleus and is substituted by ARS2c. ARS2c enhances the binding of UPF1 to NCBP1, ERF1 and DHX34, promoting the formation of the SURF complex. The presence of an EJC more than 50 nt downstream of the termination codon favors the formation of the DECID complex and transcript degradation though NMD. Left panel: ARS2n and ARS2c interactors reported in our study and/or the literature.

BioID is a widely used method that allows the detection of transient, weak and temporally separated interactions^35,50^. For this reason, it is uniquely suited to detect ARS2 isoforms interactomes, since ARS2 is known to function at multiple stages in RNA metabolism promoting the formation of highly dynamic, transient, and mutually exclusive RNP complexes. ARS2n interactome returned previously known associations with splicing factors, EJC, PHAX and the export machinery, which provided an important validation of our approach. Unexpectedly, we did not find significant enrichment of NCBP1 or NCBP2 in our data. We verified that NCBP2 was present in BioID2-ARS2n-3flag immunoprecipitations, confirming that the BioID2 tag did not affect ARS2n binding to NCBP1/2. Interestingly, we found a significant enrichment of NCBP3 in the ARS2n data. NCBP3 has been shown to interact with CBC-ARS2 and the EJC post-splicing, and functions to promote multiexonic polyadenylated mRNA export^5,46^. The presence of NCBP3, as well as PHAX, in this dataset suggests that the inability to detect NCBP1 and NCBP2 may be due to steric constraints from the placement of the biotin ligase at the N-terminus. Thus, consistent with other studies, our data illustrates that BioID is a complementary approach to AP-MS and does not replace it.

We identified the NMD pathway as one of the enriched biological processes that is shared between ARS2n and ARS2c interactomes. Using a tethering assay, NMD reporters, RT-qPCR of endogenous targets and both gain-of-function and loss-of-function experiments, we found that ARS2n inhibits while ARS2c promotes NMD. The role of ARS2n appears to be primarily in the nucleus through its interactions with the EJC and export machinery, whereas ARS2c promotes the interactions between UPF1 and NCBP1/ERF1/DHX34/SMG7 in the cytoplasm (Fig. 6E). To directly test whether the isoforms are functionally the same and differ only in cellular localization, or whether the isoforms are functionally distinct and differ in location and function, we created a cytoplasmic version of ARS2n in which we deleted the NLS but left the N-terminal leg and remaining unstructured N-terminus intact. We show that deletion of the nuclear localization signal is sufficient to confer cytoplasmic localization to the full-length isoform and abrogates the nuclear isoform’s ability to inhibit NMD (Fig. 6A, B). However, despite the cytoplasmic localization, the ΔNLS mutant was unable to promote the association of UPF1 with ERF1 and NCBP1 (Fig. 6E). This result indicates that cytoplasmic expression of ARS2n is not sufficient, and that the isoforms are functionally different.

The functional distinctness of the isoforms must be related to the differences in their N-termini. The N-terminal leg of ARS2n physically associates with the C-terminus of the protein in the crystal structure and is thus poised to affect binding events at the C-terminus^20^. Consistent with this prediction, we have shown previously that mutations within the N-terminal leg alter the RNA binding ability at the C-terminal leg. Specifically, mutation of three conserved tyrosine sites within the N-terminal leg reduces the association of ARS2 with RNA^26^. Interestingly, two of these tyrosine residues are also phosphorylated within the phosphoproteome data set, suggesting interactions to this region of the protein may be regulated^51,52^. It is possible that the presence of the N-terminal leg sterically hinders ARS2n from promoting NMD and that an isoform switch is necessary for the pathway. However, understanding how these isoforms precisely work to coordinate mRNA fate during NMD will require further study.

ARS2 switching also appears to be important for the proteomic remodelling and cell sensitivity in response to arsenic treatment. Our findings show that ARS2c is upregulated in response to arsenic while ARS2n is downregulated. Both proteomic remodeling and arsenic sensitivity were lost upon ARS2c knockdown. Moreover, while specific knockdown of ARS2c strongly increased resistance to arsenic, knockdown of both isoforms decreased this resistance, suggesting ARS2n may confer resistance to arsenic. Interestingly, ARS2c mediated arsenic sensitivity appears to be independent of the CBC complex. NCBP1 expression, like UPF1 or ARS2n, decreased following arsenic treatment, and depletion of NCBP1 had no effect on the survival of arsenic treated cells. Furthermore, co-depletion of ARS2c and NCBP1 had a similar phenotype to depletion of ARS2c alone. ARS2c mediated arsenic sensitivity is, to our knowledge, the first described function for mammalian ARS2 that is potentially independent from the CBC complex.

How ARS2c participates in arsenic sensitivity and proteomic remodeling is unclear. The mechanism appears to be unrelated to the effect of arsenic on translation inhibition. Using puromycin to inhibit translation, we found that although puromycin represses expression of ARS2n and induces expression of ARS2c, knockdown of ARS2c had no effect on the survival of puromycin treated cells (Suppl. Fig. 5G). It is also unlikely that the sensitivity to arsenic conferred by ARS2c is due to its role in NMD, since both UPF1 and NCBP1 are downregulated in response to arsenic treatment (Fig. 3B, C). Mechanistically, arsenic generates genotoxic reactive oxygen species, ER stress, promotes the induction of the unfolded protein response (UPR) and triggers p53-regulated mitochondrial apoptosis^53^. Interestingly, our BioID data showed interaction between ARS2c and proteins involved in cell redox homeostasis, protein folding, and response to unfolded protein (Fig. 4A). How such interactions regulate the cellular response to arsenic will need to be addressed in the future.

Our study shows that in high *ARS2* expressing tumours, there is an enrichment of genes clustered in cell cycle-related processes such as cell cycle DNA replication, chromosome segregation and meiotic cell cycle progress, which is consistent with the role of ARS2 during cell cycle progression^26,54^. Strikingly, the NMD pathway was strongly enriched in almost all analyzed tumours, which suggests that ARS2 mediated NMD regulation has consequences *in vivo*. The relationship between NMD and cancer development is well supported in the literature^49^. For example, in pancreatic adenosquamous carcinoma, downregulation of NMD through mutation of UPF1 promotes tumorigenesis by allowing the expression of a dominant negative PTC-bearing p53 tumour suppressor gene^24,49^. Interestingly, our data shows that high expression of *ARS2* in pancreatic cancer is associated with NMD and patient survival, which suggests that NMD regulation by ARS2 could have clinical implications in pancreatic tumours.

Understanding the role of ARS2 isoforms in cancer will require *ARS2c* specific sequence data which is not currently annotated in existing data sets. Although we could not differentiate between ARS2 isoforms in our analysis, a recent study in glioblastoma that used western blotting to detect ARS2 expression in patient tumor samples, showed expression of both ARS2 isoforms in advanced stages of the disease (see Fig.1F and supplemental Figure with original blots)^55^. Detection of specific *ARS2* isoform expression in tumour samples by RNAseq will be challenging because of the strong similarity of *ARS2* isoforms sequences (differ only by retention of intron 5) and due to the variability of *ARS2* isoforms expression, which will likely differ between cancer type, patients, and stage^55^.To address these problems, targeted methods that focus on identifying ARS2c transcripts will be needed. However, these efforts will likely be important for prognosis and potential treatments. A recent study in pancreatic cell lines showed that arsenic trioxide (ATO) induces apoptosis *in vitro* and tumour reduction in nude mice^23^. Although ATO has been effectively used in the treatment of acute promyelocytic leukemia (APL)^21,56,57^, the use of arsenic as an anticancer agent is limited by its toxicity and the lack of mechanistic studies in different tumour scenarios. Our study shows that ARS2c expression is critical for arsenic sensitivity. Therefore, ARS2c expression levels within tumours could serve as a prognostic factor to stratify those patients most likely to benefit from ATO treatment. Understanding the mechanism of ARS2c-mediated arsenic sensitivity could lead to new treatment options that promote higher arsenic toxicity within tumours.

## Supporting information

Supplemental sequences

## Data availability

All codes used in this study are publicly available. The mass spectrometry proteomics data have been deposited to the ProteomeXchange Consortium via the PRIDE^58^ partner repository.

## Supplementary data

Supplementary data are available at NAR online.

## Acknowledgements

We thank Darryl Hardie and Jason Serpa and the staff of the UVIC Genome BC Proteomics Centre for the support in the LC/MS-MS data acquisition and analysis. We thank Ryan Erdman and Scott Scholz from BCMB department Biotech Support for equipment assistance. pNMD+ and pNMD-reporter plasmids were kindly provided by K. Lukyanov ^28^. RNT1-GFP was a gift from H. Dietz (Addgene plasmid # 17708) ^29^. pAc5.1C-FLuc-Stop-5BoxB was a gift from Elisa Izaurralde (Addgene plasmid # 21301) ^30^. CBP20-3flag was a gift from Torben Heick Jensen and John LaCava ^31^. Anti-ARS2 (XL12.2 and LX186.3) was generously provided by the Ludwig Institute for Cancer Research. The results presented in Fig.7 are based upon data generated by the TCGA Research Network: https://www.cancer.gov/tcga.

