## Supplemental sequences for "Cytoplasmic Switch of ARS2 Isoforms Promotes Nonsense-Mediated mRNA Decay and Arsenic Sensitivity"

1) **ARS2n (AF/AR)**

AGAGCGCAGCGATTATGACCGTTTCCCCGGGAAAGGGATGAAAGACGGCGAGGG
GACGATTGGAATGACCGAGAGTGGGACCGTGGCCGGGAGCGCCGCAGTCGGGGT
GAATATCGAGACTACGACAGGAACCGAAGGGAACGCTTCTCTCCCCCTCGACACG
AACTCAGCCCCCCCCAGAAGCGCATGCGGAGAGACTGGGATGAGCACAGCTCTGA
CCCATACCACAGTGGCTATGACATGCCCTATGCTGGGGGGGGTGGGGGACCAACT
TACGGCCCCCCTCAGCCCTGGGGCCACCCAGACGTCCACATCATGCAGCACCATG
TCCTGCCCATCCAGGCCAGGCTGGGCAGCATCGCAGAGATTGACTTGGGGGTGCC
ACCGCCCATAATGAAGTCCTTCAAAGAGTTCCTCCTGTCTCTGGATGACTCTGTGG
ATGAGACGGAGGCAGTTAAACGCTACCATGACTACAAGCTGGACTTCCGAAGGCA
GCAGATGCAGGACTTTTTCCTGGCTCACAAAGACGAGGAGTGGTTCCGATCTAAGT
ACCACCCTGATGAGGTGGGAAAGCGTCGGCAGGAGGCCCGGGGGGCCCTGCAGA
ACCGGCTGAAGGTGTTCCTGTCCCTCATGGAGAGTGGCTGGTTTGATAACCTTCTC
TTGGACATAGACAAAGCTGATGCCATTGTCAAGATGCTAGATGCAGCTGTCATTAA
GATGGAAGGTGGCACAGAGAACGATCTCCGAATTTTGGAGCAGGAGGAGGAGGAG
GAACAGGCAGGCAAGACTGGGGAGGCCAGCAAGAAAGAGGAGGCCCGTGCTGGA
CCAGCCCTGGGGGAAGGAGAGCGCAAAGCCAATGATAAGGATGAGAAGAAAGAAG
ATGGAAAACAGGCTGAGAATGACAGTTCCAACGATGACAAAACTAAAAAATCTGAG
GGTGATGGGGACAAGGAGGAGAAGAAAGAAGAGGCTGAGAAAGGAAGCCAAAAA
GAGCAAGAAGCGGAACAGGAAGCAGAGTGGCGATGACAGCTTCGATGAGGNAGNT
GTGTCA

2) **Intron 5 region (1618-3328) based on transcript XM_030254927.2**

**2.1) Primer P1:**

X4 F:
NNNNNNNNNNNNTNNNTGGTTTNNNTGGAGGAGCTGCAGCCCTGTCCTCTGGTTC
TCCCCACCGTGGCACTTCCTGGGAGTGGGGAAGGGGAGGGAAACACTGTTGGCT
GGTGGGATCACACCCACCTCCTTCTCTGCCTTCCTCACTGGGACACCTCCAGGCA
GCTGACCAGAGCCACTTAGCCTCTTGGTACAGCACACAGACTGCGATCCTCTTGTT
GGTGTGGGTCATTGTCATCCCCAGTCAACAGGACCCAGTGTGGCCGTGGTCTCCC
AAAGGCAACGAGTGAAGCCAGACAAGAACAGCCAAGATNTANAGGGGANCCGTTT
A

X4 R:
NNNNNNNNNNNNTNGTTGCCTTATGGGANANCACGGCCACACTGGGTCCTGTTGA
CTGGGGATGACAATGACCCACACCAACAAGAGGATCGCAGTCTGTGTGCTGTACC
AAGAGGCTAAGTGGCTCTGGTCAGCTGCCTGGAGGTGTCCCAGTGAGGAAGGCAG
AGAAGGAGGTGGGTGTGATCCCACCAGCCAACAGTGTTTCCCTCCCCTTCCCCACT
CCCAGGAAGTGCCACGGTGGGGAGAACCAGAGGACAGGGCTGCAGCTCCTCCAG
CCAAACCAGAGAAATGGGCCAGGGCCAAGGGGGCAGGCANNAGGGGGGCACTAA
A

**Sequence: 100% homology transcript variants XM_030254927.2, XM_030254928.2 and XM_036165632.1 (1672-1959 from XM_030254927.2)**

TGGAGGAGCTGCAGCCCTGTCCTCTGGTTCTCCCCACCGTGGCACTTCCTGGGAG
TGGGGAAGGGGAGGGAAACACTGTTGGCTGGTGGGATCACACCCACCTCCTTCTC
TGCCTTCCTCACTGGGACACCTCCAGGCAGCTGACCAGAGCCACTTAGCCTCTTG
GTACAGCACACAGACTGCGATCCTCTTGTTGGTGTGGGTCATTGTCATCCCCAGTC
AACAGGACCCAGTGTGGCCGTGGTCTCCCAAAGGCAACGAGTGAAGCCAGACAAG
AACAGCCAAGAT

**2.2) Primer P5:**

X3sp2 F:
NNNNNNNNNNNNNNNNNNNNNNTGTTCTTGCTCTCTGCTACACACTTCTTGTTTCT
GTTTTCTCAACCTGTCTGTCTTTGTTTTAGATTTCGGGGCCCTCACATTTTCCCCATT
TCTCCTTTTCTCCTACCCCTAATTTCTTCTGCCTTTGTTTTTCTCCCCTGCCCTCAGA
GGCCTATTAAAATGTTTTCTTTTTCATGCTCTGCCCAGATCCTAACCCTCTAGCCCT
CTTCTATCCCATGGCCTACCTGAGGCTGTGCAGGCACTTGGCCTCTTTCTTTGCTC
TTTCCTGCCCCTCTGCCTTTTCTTCCTTCGAGGGGAGAAAGGTGTCCTGCCTTTTG
GCCTTTTCTATCTGGTACACTTGAGGTTTCCTTCTGGCTCCACCTGGGGTGGAGGG
AGTTGGTGGGAACCTCATCTGGGCTTGGCCAGGAGAGGGAGGTTCTCTGGGGCTC
CTGCTCCCTCAGCTTTTCTGAAGCTATGGGAAAGGGCACCTTCTCAGCCTAGTGCT
TCCCAGAGGCTCACAGCAGTAGAGCTGCCTCATAACCCTGGCCTTCCCCAGAGCC
ATGGTCCTCCTAACCCAGAGCTCTCAATTCCTGGTTCTCCCCAGATCTTTCCAGAGT
ACGACCTCTGGTTCCTTTTGGGGTGTTGGGTGGTTTTAGTGCACCCCTTTCTTGATT
TTAAGACCTTCTGTGGGTATGAGAGGCTATGCCCTGCCTTTCTTGGGCCCCTCCCC
ACATTCTGGTCTTGACTGCTCGTAACTGTTTTGTCTCTCCCATCCTGCTTGCTTTCTT
GTCCAGGTTCCGATCTAAGTACCACCCTGATGAGGTGGGAAAGCGTCGGCAGGAG
GCCCGGGGGGCCCTGCAGAACCGGCTGAAGGTGTTCCTGTCCCTCATGGAGAGT
GGCTGGTTTGATACCTTCTCTGGACTAGANAAGCTGATGCCTTGTCAGATGCTAAA
GCAGCTGTCTTAGNGGAAGGNGGCCAAAAACGATNTCCATTTGGAACNGGGGAGG
NGANAANNGCGGCAAATGGGAGCNACAAAAANNNCCGGCTGAACCCNGGGGAGG
AAGCCACCCNNANGGAAAAAAAAAGGAAGGGGNGNNTCCCGGAAANTTTTTGGGG
GGGNNNAAAAAAAAAAAANCANGNAAANNCCACTCCCCCTTTTTNNTNNNNNNNNNN
NTCTCGNGAGGGGNNNNN

X3p3 R:
NNNNNNNNNNNNNTTCTGTTCCGCTTCTTGCTCTTTTTGGCTTCCTTCTCAGCCTCTT
CTTTCTTCTCCTCCTTGTCCCCATCACCCTCAGATTTTTTAGTTTTGTCATCGTTGGA
ACTGTCATTCTCAGCCTGTTTTCCATCTTCTTTCTTCTCATCCTTATCATTGGCTTTG
CGCTCTCCTTCCCCCAGGGCTGGTCCAGCACGGGCCTCCTCTTTCTTGCTGGCCT
CCCCAGTCTTGCCTGCCTGTTCCTCCTCCTCCTCCTGCTCCAAAATTCGGAGATCG
TTCTCTGTGCCACCTTCCATCTTAATGACAGCTGCATCTAGCATCTTGACAATGGCA
TCAGCTTTGTCTATGTCCAAGAGAAGGTTATCAAACCAGCCACTCTCCATGAGGGA
CAGGAACACCTTCAGCCGGTTCTGCAGGGCCCCCCGGGCCTCCTGCCGACGCTTT
CCCACCTCATCAGGGTGGTACTTAGATCGGAACCTGGACAAGAAAGCAAGCAGGA
TGGGAGAGACAAAACAGTTACGAGCAGTCAAGACCAGAATGTGGGGAGGGGCCCA
AGAAAGGCAGGGCATAGCCTCTCATACCCACAGAAGGTCTTAAAATCAAGAAAGGG
GTGCACTAAAACCACCCAACACCCCAAAAGGAACCAGAGGTCGTACTCTGGAAAGA
TCTGGGGAGAACCAGGAATTGAGAGCTCTGGGTTAGGAGGACCATGGCTCTGGAG
AAGGCCAGGGTTATGAGGCAGCTCTACTGCTGTGAGCCTCTGGGAAGCACTAGGC

TGAGAAGGTGCCCTTTCGCATAGCTTCAGAAAAGCTGAGGGAGCAGGAGCCCCAG
AGAACCTCCCTCTCCTGGGCAAGCCCAGATGAGGTTCCACCAACTNCCTCCACCCA
GGTGGNNNCGAAGGAAACCTCAGTGTACCAANNNNAAAGGCAAAAGGNNGAAACT
TTCTCCCTNGAAGAAAAAAGGNANGGGCAGAAAAACCAAAA

**Sequence: 99% homology transcript variants XM_030254927.2, XM_006504631.3 and XM_036165632.1 (2554- 3799 from XM_030254927.2)**

TGTTCTTGCTCTCTGCTACACACTTCTTGTTTCTGTTTTCTCAACCTGTCTGTCTTTG
TTTTAGATTTCGGGGCCCTCACATTTTCCCCATTTCTCCTTTTCTCCTACCCCTAATT
TCTTCTGCCTTTGTTTTTCTCCCCTGCCCTCAGAGGCCTATTAAAATGTTTTCTTTTT
CATGCTCTGCCCAGATCCTAACCCTCTAGCCCTCTTCTATCCCATGGCCTACCTGA
GGCTGTGCAGGCACTTGGCCTCTTTCTTTGCTCTTTCCTGCCCCTCTGCCTTTTCTT
CCTTCGAGGGGAGAAAGGTGTCCTGCCTTTTGGCCTTTTCTATCTGGTACACTTGA
GGTTTCCTTCTGGCTCCACCTGGGGTGGAGGGAGTTGGTGGGAACCTCATCTGGG
CTTGGCCAGGAGAGGGAGGTTCTCTGGGGCTCCTGCTCCCTCAGCTTTTCTGAAG
CTATGGGAAAGGGCACCTTCTCAGCCTAGTGCTTCCCAGAGGCTCACAGCAGTAG
AGCTGCCTCATAACCCTGGCCTTCCCCAGAGCCATGGTCCTCCTAACCCAGAGCTC
TCAATTCCTGGTTCTCCCCAGATCTTTCCAGAGTACGACCTCTGGTTCCTTTTGGGG
TGTTGGGTGGTTTTAGTGCACCCCTTTCTTGATTTTAAGACCTTCTGTGGGTATGAG
AGGCTATGCCCTGCCTTTCTTGGGCCCCTCCCCACATTCTGGTCTTGACTGCTCGT
AACTGTTTTGTCTCTCCCATCCTGCTTGCTTTCTTGTCCAGGTTCCGATCTAAGTAC
CACCCTGATGAGGTGGGAAAGCGTCGGCAGGAGGCCCGGGGGGCCCTGCAGAAC
CGGCTGAAGGTGTTCCTGTCCCTCATGGAGAGTGGCTGGTTTGATAACCTTCTCTT
GGACATAGACAAAGCTGATGCCATTGTCAAGATGCTAGATGCAGCTGTCATTAAGA
TGGAAGGTGGCACAGAGAACGATCTCCGAATTTTGGAGCAGGAGGAGGAGGAGGA
ACAGGCAGGCAAGACTGGGGAGGCCAGCAAGAAAGAGGAGGCCCGTGCTGGACC
AGCCCTGGGGGAAGGAGAGCGCAAAGCCAATGATAAGGATGAGAAGAAAGAAGAT
GGAAAACAGGCTGAGAATGACAGTTCCAACGATGACAAAACTAAAAAATCTGAGGG
TGATGGGGACAAGGAGGAGAAGAAAGAAGAGGCTGAGAAGGAAGCCAAAAAGAGC
AAGAAGCGGAACAG

**2.3) Primer P3**

X3sp2 F:
NNNNNNNNTNNTTNTCTNTNNTGTTCTTGCTCTCTGCTACACACTTCTTGTTTCTGTT
TTCTCAACCTGTCTGTCTTTGTTTTAGATTTCGGGGCCCTCACATTTTCCCCATTTCT
CCTTTTCTCCTACCCCTAATTTCTTCTGCCTTTGTTTTTCTCCCCTGCCCTCAGAGGC
CTATTAAAATGTTTTCTTTTTCATGCTCTGCCCAGATCCTAACCCTCTAGCCCTCTTC
TATCCCATGGCCTACCTGAGGCTGTGCAGGCACTTGGCCTCTTTCTTTGCTCTTTC
CTGCCCCTCTGCCTTTTCTTCCTTCGAGGGGAGAAAGGTGTCCTGCCTTTTGGCCT
TTTCTATCTGGTACACTTGAGGTTTCCTTCTGGCTCCACCTGGGGTGGAGGGAGTT
GGTGGGAACCTCATCTGGGCTTGGCCAGGAGAGGGAGGTTCTCTGGGGCTCCTG
CTCCCTCAGCTTTTCTGAAGCTATGGGAAAGGGCACCTTCTCAGCCTAGTGCTTCC
CAGAGGCTCACAGCAGTAGAGCTGCCTCATAACCCTGGCCTTCCCCAGAGCCATG
GTCCTCCTAACCCAGAGCTCTCAATTCCTGGTTCTCCCCAGATCTTTCCAGAGTAC
GACCTCTGGTTCCTTTTGGGGTGTTGGGTGGTTTTAGTGCACCCCTTTCTTGATTTT
AAGACCTTCTGTGGGTATGAGAGGCTATGCTNTGNNTTTTTTTGGGGGNGAAAGC

X3sp2 R:
NNNNNNNNNNNNNNNTTNNNTNNAGAAAGGGGTGCACTAAAACCACCCAACACCCC
AAAAGGAACCAGAGGTCGTACTCTGGAAAGATCTGGGGAGAACCAGGAATTGAGA
GCTCTGGGTTAGGAGGACCATGGCTCTGGGGAAGGCCAGGGTTATGAGGCAGCTC
TACTGCTGTGAGCCTCTGGGAAGCACTAGGCTGAGAAGGTGCCCTTTCCCATAGCT
TCNGAAAAGCTGAGGGAGCAGGAGCCCCNGANAACCTCCCTCTCCTGGNCNAGCC
CNGATGANGTTNCCNCCNACTNCCTNCNCCCCNGGTGG

**Sequence: 99% homology transcript variants XM_030254927.2, XM_006504631.3 and XM_036165632.1 (2554- 3240 from XM_030254927.2)**
TGTTCTTGCTCTCTGCTACACACTTCTTGTTTCTGTTTTCTCAACCTGTCTGTCTTTG
TTTTAGATTTCGGGGCCCTCACATTTTCCCCATTTCTCCTTTTCTCCTACCCCTAATT
TCTTCTGCCTTTGTTTTTCTCCCCTGCCCTCAGAGGCCTATTAAAATGTTTTCTTTTT
CATGCTCTGCCCAGATCCTAACCCTCTAGCCCTCTTCTATCCCATGGCCTACCTGA
GGCTGTGCAGGCACTTGGCCTCTTTCTTTGCTCTTTCCTGCCCCTCTGCCTTTTCTT
CCTTCGAGGGGAGAAAGGTGTCCTGCCTTTTGGCCTTTTCTATCTGGTACACTTGA
GGTTTCCTTCTGGCTCCACCTGGGGTGGAGGGAGTTGGTGGGAACCTCATCTGGG
CTTGGCCAGGAGAGGGAGGTTCTCTGGGGCTCCTGCTCCCTCAGCTTTTCTGAAG
CTATGGGAAAGGGCACCTTCTCAGCCTAGTGCTTCCCAGAGGCTCACAGCAGTAG
AGCTGCCTCATAACCCTGGCCTTCCCCAGAGCCATGGTCCTCCTAACCCAGAGCTC
TCAATTCCTGGTTCTCCCCAGATCTTTCCAGAGTACGACCTCTGGTTCCTTTTGGGG
TGTTGGGTGGTTTTAGTGCACCCCTTTCTTGATTTTAAGACCTTCTGTGGGTATGAG
AGGCTATGCT

**2.4) Primer P4**

X3fl F:
NNNNNTNNNNNNANNANGTTCTCTGGGGCTCCTGCTCCCTCAGCTTTTCTGAAGCT
ATGGGAAAGGGCACCTTCTCAGCCTAGTGCTTCCCAGAGGCTCACAGCAGTAGAG
CTGCCTCATAACCCTGGCCTTCCCCAGAGCCATGGTCCTCCTAACCCAGAGCTCTC
AATTCCTGGTTCTCCCCAGATCTTTCCAGAGTACGACCTCTGGTTCCTTTTGGGGTG
TTGGGTGGTTTTAGTGCACCCCTTTCTTGATTTTAAGACCTTCTGTGGGTATGAGAG
GCTATGCCCTGCCTTTCTTGGGCCCCTCCCCACATTCTGGTCTTGACTGCTCGTAA
CTGTTTTGTCTCTCCCATCCTGCTTGCTTTCTTGTCCAGGTTCCGATCTAAGTACCA
CCCTGATGAGGTGGGAAAGCGTCGGCAGGAGGCCCGGGGGGCCCTGCAGAACCG
GCTGAAGGTGTTCCTGTCCCTCATGGAGAGTGGCTGGTTTGATAACCTTCTCTTGG
ACATAGACAAAGCTGATGCCATTGTCAAGATGCTAGATGCAGCTGTCATTAAGATG
GAAGGTGGCACAGAGAACGATCTCCGAATTTTGGAGCAGGAGGAGGAGGAGGAAC
AGGCAGGCAAGACTGGGGAGGCCAGCAAGAAAGAGGAGGCCCGTGCTGGACCAG
CCCTGGGGGAAGGAGAGCGCAAAGCCCATGATAAGGATGAGAAGAAAGAAGATGG
AAAACAGGCTGAGAATGACAGTTCCCACGATGACAAAACTAAAAATCTGAGGGTGA
TGGGGACAGGAGGAGAAAAAAGANNGGCTGAGAGGAGCCCCAAGAGCAGAACCG
GANAGGAGCCAAATGCGAGNCNNT

X3p3 R:
NNNNNNNNTCTGNTTNNTGTTCCGCTTCTTGCTCTTTTTGGCTTCCTTCTCAGCCTC

TTCTTTCTTCTCCTCCTTGTCCCCATCACCCTCAGATTTTTTAGTTTTGTCATCGTTG
GAACTGTCATTCTCAGCCTGTTTTCCATCTTCTTTCTTCTCATCCTTATCATTGGCTT
TGCGCTCTCCTTCCCCCAGGGCTGGTCCAGCACGGGCCTCCTCTTTCTTGCTGGC
CTCCCCAGTCTTGCCTGCCTGTTCCTCCTCCTCCTGCTCCAAAATTCGGAGAT
CGTTCTCTGTGCCACCTTCCATCTTAATGACAGCTGCATCTAGCATCTTGACAATGG
CATCAGCTTTGTCTATGTCCAAGAGAAGGTTATCAAACCAGCCACTCTCCATGAGG
GACAGGAACACCTTCAGCCGGTTCTGCAGGGCCCCCCGGGCCTCCTGCCGACGC
TTTCCCACCTCATCAGGGTGGTACTTAGATCGGAACCTGGACAAGAAAGCAAGCAG
GATGGGAGAGACAAAACAGTTACGAGCAGTCAAGACCAGAATGTGGGGAGGGGCC
CAAGAAAGGCAGGGCATAGCCTCTCATACCCACAGAAGGTCTTAAAATCAAGAAAG
GGGTGCACTAAAACCACCCAACACCCCAAAAGGAACCAGAGGTCGTACTCTGGAA
AGATCTGGGGAGAACCAGGAATTGAGAGCTCTGGGTTAGGAGGACCATGGCTCTG
GGGAAGGCCAGGGTTATGAGGCAGCTCTACTGCTGTGAGCCTCTGGGAAGCACTA
GGCTGAGAAGGTGCCCTTTCCCATAGCTTCAGAAAGCGAGGGGAGCAGGAGCCCC
ACAGAACTCCCTCTCTGGCAGCCATGAGTTTCCCCCAAATCACAAAAAAAAAAACAC
NAAAAGGGGGAGGGGCCCTTTCCCTCCCNTCAAAAAGCTGAGGGANNNAGGAGCC
CCAGAAAACTCCCNCTCCTGGCCAACCCAAAGAAAGTTCCCCCACCCNACTACCCC
AANNNAGGGAAGGGANANCCCAAAACCCCCCNCCCGG

**Sequence: 99% homology transcript variants XM_030254927.2, XM_006504631.3 and XM_036165632.1 (2970- 3798 from XM_030254927.2)**
GTTCTCTGGGGCTCCTGCTCCCTCAGCTTTTCTGAAGCTATGGAAAGGGCACCTT
CTCAGCCTAGTGCTTCCCAGAGGCTCACAGCAGTAGAGCTGCCTCATAACCCTGG
CCTTCCCCAGAGCCATGGTCCTCCTAACCCAGAGCTCTCAATTCCTGGTTCTCCCC
AGATCTTTCCAGAGTACGACCTCTGGTTCCTTTTGGGGTGTTGGGTGGTTTTAGTG
CACCCCTTTCTTGATTTTAAGACCTTCTGTGGGTATGAGAGGCTATGCCCTGCCTTT
CTTGGGCCCCTCCCCACATTCTGGTCTTGACTGCTCGTAACTGTTTTGTCTCTCCCA
TCCTGCTTGCTTTCTTGTCCAGGTTCCGATCTAAGTACCACCCTGATGAGGTGGGA
AAGCGTCGGCAGGAGGCCCGGGGGGCCCTGCAGAACCGGCTGAAGGTGTTCCTG
TCCCTCATGGAGAGTGGCTGGTTTGATAACCTTCTCTTGGACATAGACAAAGCTGA
TGCCATTGTCAAGATGCTAGATGCAGCTGTCATTAAGATGGAAGGTGGCACAGAGA
ACGATCTCCGAATTTTGGAGCAGGAGGAGGAGGAGGAACAGGCAGGCAAGACTGG
GGAGGCCAGCAAGAAAGAGGAGGCCCGTGCTGGACCAGCCCTGGGGGAAGGAGA
GCGCAAAGCCAATGATAAGGATGAGAAGAAAGAAGATGGAAAACAGGCTGAGAAT
GACAGTTCCAACGATGACAAAACTAAAAAATCTGAGGGTGATGGGGACAAGGAGGA
GAAGAAAGAAGAGGCTGAGAAGGAAGCCAAAAAGAGCAAGAAGCGGAACA

3) **Full predicted sequence amplification (XF/3UTR)**
Primer3 F:
NNNGNNNNNNNNNNNNNNNNNNNNNGCTCGTACTGTTTTGTCTCTCCCATCCTGCTT
GCTTTCTTGTCCAGGTTCCGATCTAAGTACCACCCTGATGAGGTGGGAAAGCGTCG
GCAGGAGGCCCGGGGGGCCCTGCAGAACCGGCTGAAGGTGTTCCTGTCCCTCAT
GGAGAGTGGCTGGTTTGATAACCTTCTCTTGGACATAGACAAAGCTGATGCCATTG
TCAAGATGCTAGATGCAGCTGTCATTAAGATGGACAGGTGGCACAGAGAACGATCT
CCGAATTTTGGAGCAGGAGGCCAGGAGGAGGAACAGGCAGGCAAGACTGGGGAG
GCCAGCAAGATAGGGAGGCCCGTGCTGGACCAGCCCTGGGGGAAGGAGAGCGCA
CAGCCAATGATAAGGATGAGAAGAAAGAAGATGGAAAACAGGCTGAGAATGACAGT

TCCAACGATGACGAACTAAAAAATCTGAGGGTGATGGGGACAAGGAGGAGAAGAA
AGAAGAGGCTGAGAAGGAAGCCAAAAAGAGCAAGAAGCGGAACAGGAAGCAGAG
GGCGATGACAGCTTCGATGAGGGCAGTGTGTCCGAGTCTGAGTCCGAGTCTGAGG
ATGGCCAGGCCGAGGAGGAGAAGGAGGAGGCCGAAGACTCTTNGGNTGNANTGT
AAGCCCCNGNNCNTGNATAAGGANTTGCTCTCTCGATGCTGCCCNGGTTCGANNN
NAACCTTTCGNNGTTGCATATCATTTGCTCTCTCATNNTGCACNNCATCTTTACGCN
ANTNTCACTGTCAGAGATCCTTTCTCTTTGTGANGTATNCTCACGCTGTCTGCGAGA
GTTTTTGATCNCANCGTCANCCTTANAGGAAAGNTNTTNCGACCCTGCCTGAGNGT
CTNTTTCCNGGNAGTGTGNAACTGTNNTCGNGNTCTGATCGGNNNCTGNGANANN
NTCTNNGNNTCNNNCAGTGANTGNNNGTTNNNCNGNNNNNTGACANAAAGATGCT
GNNNCTGN

Primer4 F:
GNNNNNNNNNNNNNNNNNNNGAGGCCGAGAGCACTTAAAGAAAAGGAGAAGCCC
AAAGAGGAGGAGAAGGAGAAGCCTAAGGATGCTGCAGGGTTGGAGTGTAAGCCCC
GGCCCTTGCATAAGACTTGCTCTCTCTTCATGCGCAACATCGCACCCAACATTTCAA
GGGCAGAGATCATTTCTCTTTGTAAACGATACCCAGGCTTTATGCGAGTGGCACTG
TCAGAGCCCCAGCCAGAGAGGAGGTTTTTTCGCCGTGGCTGGGTGACTTTTGACC
GCAGTGTTAACATTAAGGAGATCTGTTGGAACCTGCAGAACATTCGGCTCCGGGAG
TGTGAACTGAGTCCCGGTGTGAACAGAGACCTGACCCGTCGTGTCCGCAACATAAA
TGGCATTACACAGCACAAGCAGATAGTGCGCAATGACATCAAGTTGGCAGCCAAGC
TAATCCACACACTGGATGACAGGACCCAGCTCTGGGCCTCTGAGCCTGGGACGCC
TCCTGTGCCCACAAGCCTCCCCTCGCAAAACCCCATCCTGAAGAACATCACTGACT
ACCTGATTGAGGAAGTGAGTGCGGAGGAGGAGGAGCTTCTGGGGAGCAGTGGGG
GACCCCCTCCTGAGGAGCCTCCCAAGGAGGGCAACCCAGCCGAGATCAATGTGGA
GAGGGATGAGAAGCTGATCAAGGTCTTGGATAAACTTCTTCTCTATTTGCGTATTGT
GCATTCTCTGGATTATTATAACACCTGTGAGTACCCTAATGAAGACGAGATGCCCAA
CCGCTGTGGCATAATCCACGTTCGGGGGCCCATGCCTCCCAACCGAATTAGTCAC
GGAGAAGTGCTGGAGTGGCAGAAGACATTTGAGGAGAAACTGACTCCACTGTTGA
GTGTGCGTGAATCCCTTTCTGAGGAAGAGGCCCANANATGGGTCGAAAAGACCCA
GAGCAGAAGTNNNAGTTGTCNCCTCCANACGCNGACTGGGCNNNTANNTGNTATG
N

Primer5 R:
NNGNNNNNNNNNNNNNNNNNNNNNNNNNNNGTCCCGATACTCCACTATGGCCCTTG
GGTCTCCTCGAACCATCCTGTTCCGAGGTTTCCCAGGATAACCGCCTTGGCCTCGA
AAAGCATCATAGTTCCCACGGCCGGCACCATATGGAGCATGGGGGTATGGAGGCC
CTCCTGTTGGGACTGCAGGGCGGACAGCACCAGCTCCATAGCCCAAGATGGGAGG
CCGGGGCTGACCATACGGCATCAAGCCCTGTGGCGTCTGATGTGGGTAGGGAAGT
CCTGGGGTCAGGCCTGGGGGGAGTATCTGGGCAGGGCCAGGTGGCTGAGCTGGC
TTGATCTCAGGCAAAGCTGGGCGCTTGGCGTCTGTGAGAAAGTTATTGAAGAACGC
CACCTCCTTCTTCACCTCCTCGATCTTCTCGGCATGCTTATTGAAGATATGCTTGCG
CACAAACTCCGGGCCCTTGAATTTCTTGACAATGACAGGACATATGCACTTATCCTT
GCCCAGTTCCTGCGTGTTGGACGTGACAAACTTCTCCACTTCTTGCTCTGGGTCTT
TTCGACCCATCTTCTGGGCCTCTTCCTTCAAAGGGATTCACGCACACTCATCAGTG
GAGTCAGTTTCTCCTCAAATGTCTTCTGCCACTCCTGCACTTCTCCGTGACTAATTC
NGTTGGCATGCANNTCCCCNTCAACGTGAATTATGTTACAGCGGTTGTGCATCTCG

TCTTCTTCANGGTACTCACAGGTGTTATACTAATCCAGAGAATGTGTCATACTGCAA
ATACAGCAGATNTTTATCCACGACTNTGATCAGCATCTCATCCCTCACCAAAGTGAA
CTCGGACTGGGNTGCACGCCCTTGGANGCTCCTCAGGAGTGNNGNTCNCNCGCT
GCTCCCANNAAGNATCATCCTCCTCAGCANTCACNTNNACAATCATGNAGTNNATG
NTGATCNTCNNCATGNGNCTNCTGCNNAGTGNANGNNNNGCNGNGNATCNN

**Sequence: 99% homology transcript variants XM_030254927.2, XM_006504631.3 and XM_036165632.1 (3282- 5336 from XM_030254927.2)**

GCTCGTACTGTTTTGTCTCTCCCATCCTGCTTGCTTTCTTGTCCAGGTTCCGATCTA
AGTACCACCCTGATGAGGTGGGAAAGCGTCGGCAGGAGGCCCGGGGGGCCCTGC
AGAACCGGCTGAAGGTGTTCCTGTCCCTCATGGAGAGTGGCTGGTTTGATAACCTT
CTCTTGGACATAGACAAAGCTGATGCCATTGTCAAGATGCTAGATGCAGCTGTCATT
AAGATGGACAGGTGGCACAGAGAACGATCTCCGAATTTTGGAGCAGGAGGCCAGG
AGGAGGAACAGGCAGGCAAGACTGGGGAGGCCAGCAAGATAGGGAGGCCCGTGC
TGGACCAGCCCTGGGGGAAGGAGAGCGCACAGCCAATGATAAGGATGAGAAGAAA
GAAGATGGAAAACAGGCTGAGAATGACAGTTCCAACGATGACGAACTAAAAAATCT
GAGGGTGATGGGGACAAGGAGGAGAAGAAAGAAGAGGCTGAGAAGGAAGCCAAA
AAGAGCAAGAAGCGGAACAGGAAGCAGAGGGCGATGACAGCTTCGATGAGGGCA
GTGTGTCCGAGTCTGAGTCCGAGTCTGAGGATGGCCAGGCCGAGGAGGAGAAGG
AGGAGGCCGAAGAAGCACTTAAAGAAAAGGAGAAGCCCAAAGAGGAGGAGAAGGA
GAAGCCTAAGGATGCTGCAGGGTTGGAGTGTAAGCCCCGGCCCTTGCATAAGACT
TGCTCTCTCTTCATGCGCAACATCGCACCCAACATTTCAAGGGCAGAGATCATTTCT
CTTTGTAAACGATACCCAGGCTTTATGCGAGTGGCACTGTCAGAGCCCCAGCCAGA
GAGGAGGTTTTTTCGCCGTGGCTGGGTGACTTTTGACCGCAGTGTTAACATTAAGG
AGATCTGTTGGAACCTGCAGAACATTCGGCTCCGGGAGTGTGAACTGAGTCCCGG
TGTGAACAGAGACCTGACCCGTCGTGTCCGCAACATAAATGGCATTACACAGCACA
AGCAGATAGTGCGCAATGACATCAAGTTGGCAGCCAAGCTAATCCACACACTGGAT
GACAGGACCCAGCTCTGGGCCTCTGAGCCTGGGACGCCTCCTGTGCCCACAAGCC
TCCCCTCGCAAAACCCCATCCTGAAGAACATCACTGACTACCTGATTGAGGAAGTG
AGTGCGGAGGAGGAGGAGCTTCTGGGGAGCAGTGGGGGACCCCCTCCTGAGGAG
CCTCCCAAGGAGGGCAACCCAGCCGAGATCAATGTGGAGAGGGATGAGAAGCTGA
TCAAGGTCTTGGATAAACTTCTTCTCTATTTGCGTATTGTGCATTCTCTGGATTATTA
TAACACCTGTGAGTACCCTAATGAAGACGAGATGCCCAACCGCTGTGGCATAATCC
ACGTTCGGGGGCCCATGCCTCCCAACCGAATTAGTCACGGAGAAGTGCTGGAGTG
GCAGAAGACATTTGAGGAGAAACTGACTCCACTGTTGAGTGTGCGTGAATCCCTTT
CTGAGGAAGAGGCCCANANATGGGTCGAAAAGACCCAGAGCAGGAAGTGGAGAA
GTTTGTCACGTCCAACACGCAGGAACTGGGCAAGGATAAGTGCATATGTCCTGTCA
TTGTCAAGAAATTCAAGGGCCCGGAGTTTGTGCGCAAGCATATCTTCAATAAGCAT
GCCGAGAAGATCGAGGAGGTGAAGAAGGAGGTGGCGTTCTTCAATAACTTTCTCAC
AGACGCCAAGCGCCCAGCTTTGCCTGAGATCAAGCCAGCTCAGCCACCTGGCCCT
GCCCAGATACTCCCCCCAGGCCTGACCCCAGGACTTCCCTACCCACATCAGACGC
CACAGGGCTTGATGCCGTATGGTCAGCCCCGGCCTCCCATCTTGGGCTATGGAGC
TGGTGCTGTCCGCCCTGCAGTCCCAACAGGAGGGCCTCCATACCCCCATGCTCCA
TATGGTGCCGGCCGTGGGAACTATGATGCTTTTCGAGGCCAAGGCGGTTATCCTG
GGAAACCTCGGAACAGGATGGTTCGAGGAGACCCAAGGGCCATAGTGGAGTATCG
GGAC

**4) Human: Intron 5 region (963-1289) based on transcript XM_024446794.1**

**4.1) Primer P1**

P1F:
CACCCGGGCTGAACGGCGGGGCCGGCCGGCAGACAGGCAGGGCACCGCTGTGC
ACAATGCAGTGGTCTAGCTGAATGTGATTGCTCAGGCAGCTAGACCACTGCATTGT
GCACAGCGGTGCCCTGCCTGTCTGCCTGGCCGCCCCCGCCGCCCAGCCCGCAGG
TATGGGCTGGGTATGGTNA

P1R:
TCACTCTGCATGAGCAATCTACCATACCCAGCCCATACATGAGCCCCCGGGCTGAG
CGGCGGGGCCGGCCAGGCAGACAGGCAGGGCACCGCTGGCACAAGNNGTCGTT

**Sequence: 93% homology transcript variant XM_024446794.1 (1027- 1148 from XM_024446794.1)**

ACCATACCCAGCCCATACATGAGCCCCCGGGCTGAGCGGCGCGGGGCCGGCCAG
GCAGACAGGCAGGGCACCGCTGTGCACAATGCAGTGGTCTAGCTGAATGTGATTG
CTCAGGCAGCTA

**4.2) Primer P2**

P2F:
GCTGACACCACCTCCACCCCCACCATACCCAGCCCATACATGAGCCCCCGGGCTG
AGCGGCGGGGCCGGCCAGGCAGACAGGCAGGGCACCGCTGTGCACAATGCAGC
GGTTTAGCTAGTGA

P2R
NNNNNNNNNTNNTGTCTGCCTGGCCGGCCCCGCCGCTCAGCCCGGGGGCTCATG
TATGGGCTGGGTATGGTGGGGGTGGAGGTGGGTTTGCTCCGAGCTCCGTCGATGA
GCGGGTGTAGTTCCCAACC

**Sequence: 95% homology transcript variant XM_024446794.1 (974- 1125 from XM_024446794.1)**

GGGAACTACACCCGCTCATCGACGGAGCTCGGAGCAAACCCACCTCCACCCCCAC
CATACCCAGCCCATACATGAGCCCCCGGGCTGAGCGGCGGGGCCGGCCAGGCAG
ACAGGCAGGGCACCGCTGTGCACAATGCAGCGGTTTAGCTAGTGA
