## Supplemental Figures for "Cytoplasmic Switch of ARS2 Isoforms Promotes Nonsense-Mediated mRNA Decay and Arsenic Sensitivity"

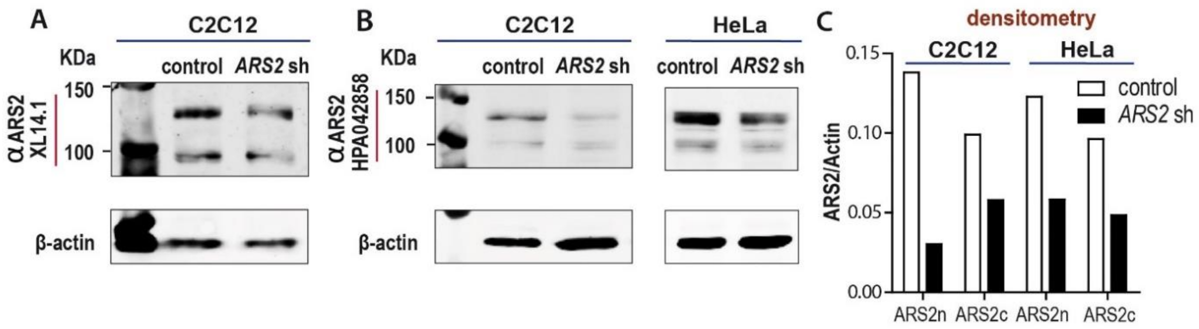


**Sup Figure 1: ARS2 antibody validation.** **A)** Validation of polyclonal antibody XL14.1. Western Blot of $C_2C_{12}$ whole cell lysate using anti-ARS2 antibody (XL14.1) showed 2 protein products. Both bands decreased when *Ars2* sh is used, since it targets all ARS2 isoforms**. B)** Western Blot of $C_2C_{12}$ and HeLa whole cell lysate using an anti-ARS2 antibody (HPA042858) validated for Western Blot by the Human Protein Atlas. https://www.proteinatlas.org/ENSG00000087087-SRRT/antibody. The image shows 2 protein products that are knocked down when *Ars2* sh is used. The results, comparable to **A)** validate XL14.1 antibody for Western Blot studies. **C)** Densitometry analysis of **B)**.

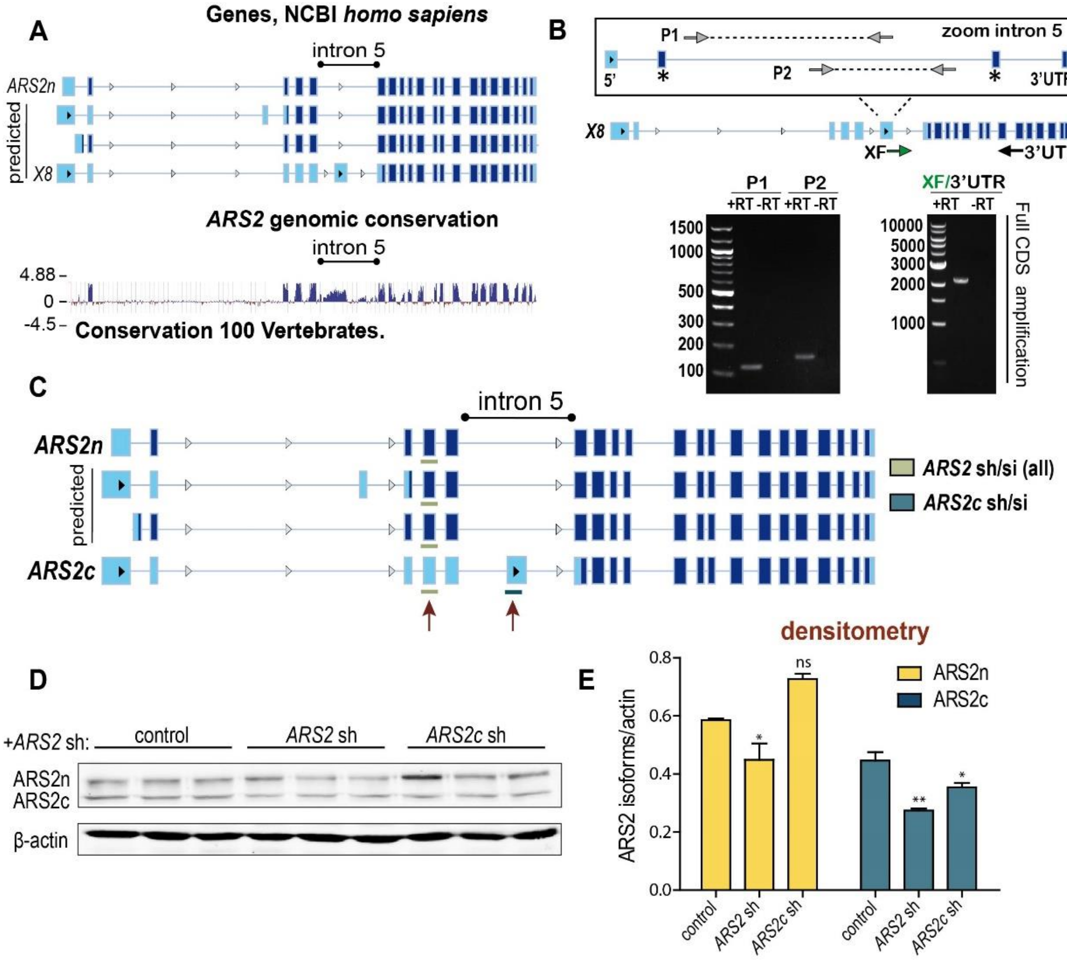


**Sup Figure 2: Cytoplasmic isoforms of *ARS2* in human cells**. **A)** ARS2 sequences reported in the NCBI Gene database (*homo sapiens*), coding sequence represented in dark blue. Bottom: conservation analysis of ARS2 between human and 100 vertebrates. The alignments were generated using Mulitz and UCSC/Penn State Bioinformatics comparative genomics alignment pipeline. Evolutionary conservation was measured using PhyloP and PhastCons. **B) (Top)** Diagram of primers used to amplify intron 5 or the full CDS of the new isoforms. **(Bottom)** RT-PCR amplifying different fragments of intron 5 (p1 and p2) or the full CDS of the isoforms (XF/3'UTR). **C)** Diagram of short hairpins used in the study for HeLa cells. *ARS2* sh targets all *ARS2* isoforms, while *ARS2*c sh is specific for the *ARS2c* isoform. **D)** Western Blot of HeLa whole cell lysate with anti-ARS2 antibody showed 2 protein products. Both bands decreased when *ARS2* sh is used, since it targets all ARS2 isoforms. *ARS2c* is specific for the ARS2c isoform, as a consequence the smaller band decreased (ARS2c) while the band on top (ARS2n) remains unaffected. β-actin is showed as a loading control. **E)** Densitometry analysis of **D).** Data are represented as mean ± SEM for n=3 biologically independent samples**.** Statistical analysis: One way ANOVA, post-test Newman-Keuls (*p≤0.05; **p≤0.01).

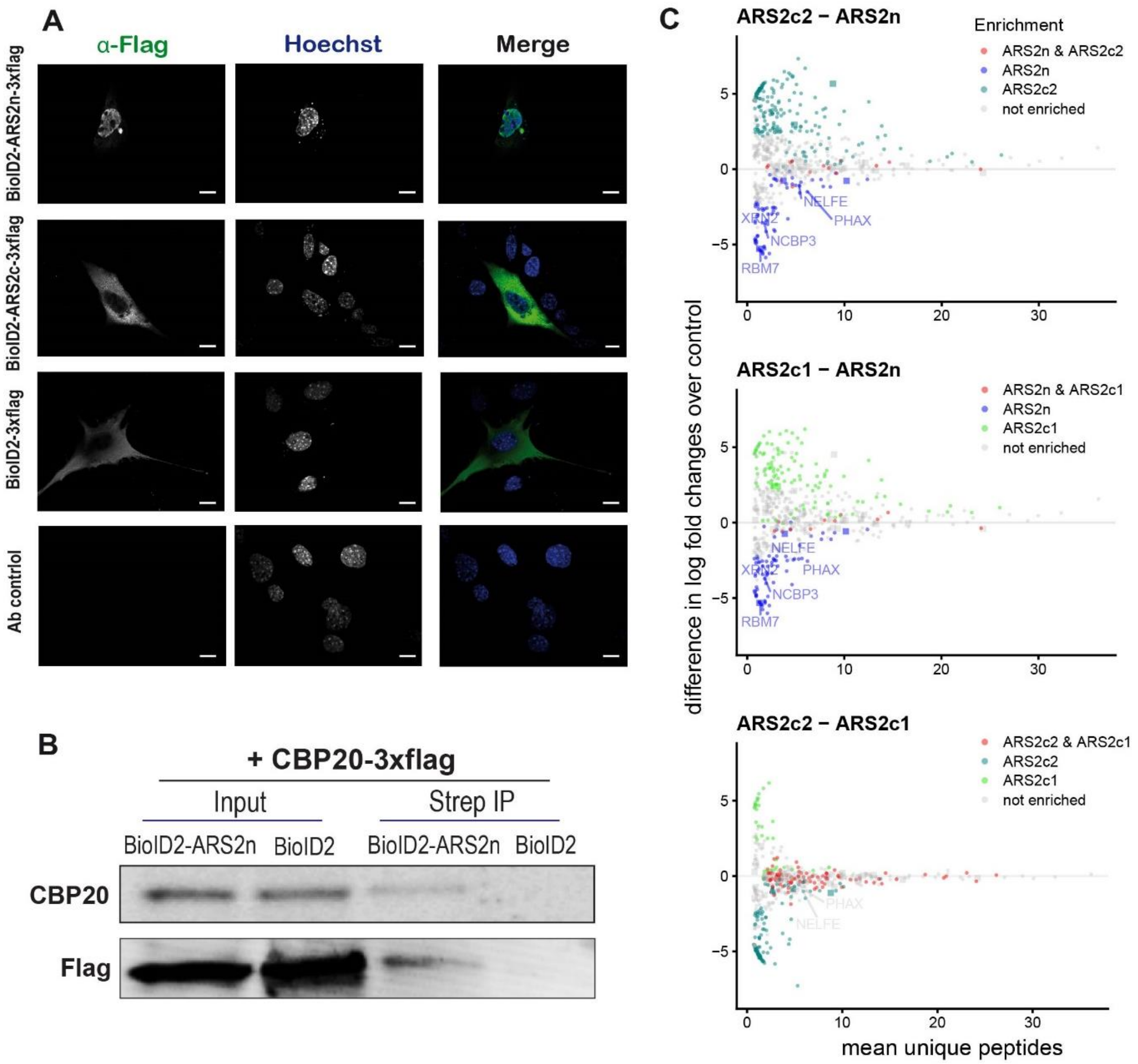


**Sup Figure 3: BioID2 tag does not affect localization or function of ARS2 isoforms. A)** HEK293T cells transfected with BioD2-ARS2n/c-3xflag showed nuclear and cytoplasmic localization, respectively.

ARS2 was detected with anti-flag antibodies (green) and nuclei were stained with Hoechst 33342 (blue). BioID2-3xflag and antibody control are also shown. Scale bar indicates 10 μm. **B)** BioID2 tag does not affect binding of ARS2 to the CBC complex. HEK293T cells were co-transfected with BioID2-ARS2n and CBP20-3xflag. After biotin treatment, lysis and streptavidin immunoprecipitation, CBP20 was detected with anti-flag and anti-CBP20 antibodies. **C)** HEK293T cells were transfected with BioID2-ARS2n/c1/c2-3xflag or control, and treated with biotin for 24h. After lysis, biotinylated interactomes of ARS2 isoforms and biotin ligase control were immunoprecipitated with streptavidin beads and detected by LC/MS-MS. 15 independenlty treated 10cm$^2$ dishes are included per condition. ARS2 isoforms interactomes were normalized against biotin ligase control and analyzed by SAINT. Only proteins with a saint score (SP>0.7) were included in the subsequent analysis. Dot blot comparison of ARS2 isoforms are shown in **C)**. ARS2n and ARS2c1/c2 do not share most of their interactors, while ARSc1 and ARS2c2 do. (matches are shown in red). Previously reported ARS2n interactors like NCBP3, PHAX and RBM7 are highlighted in blue.

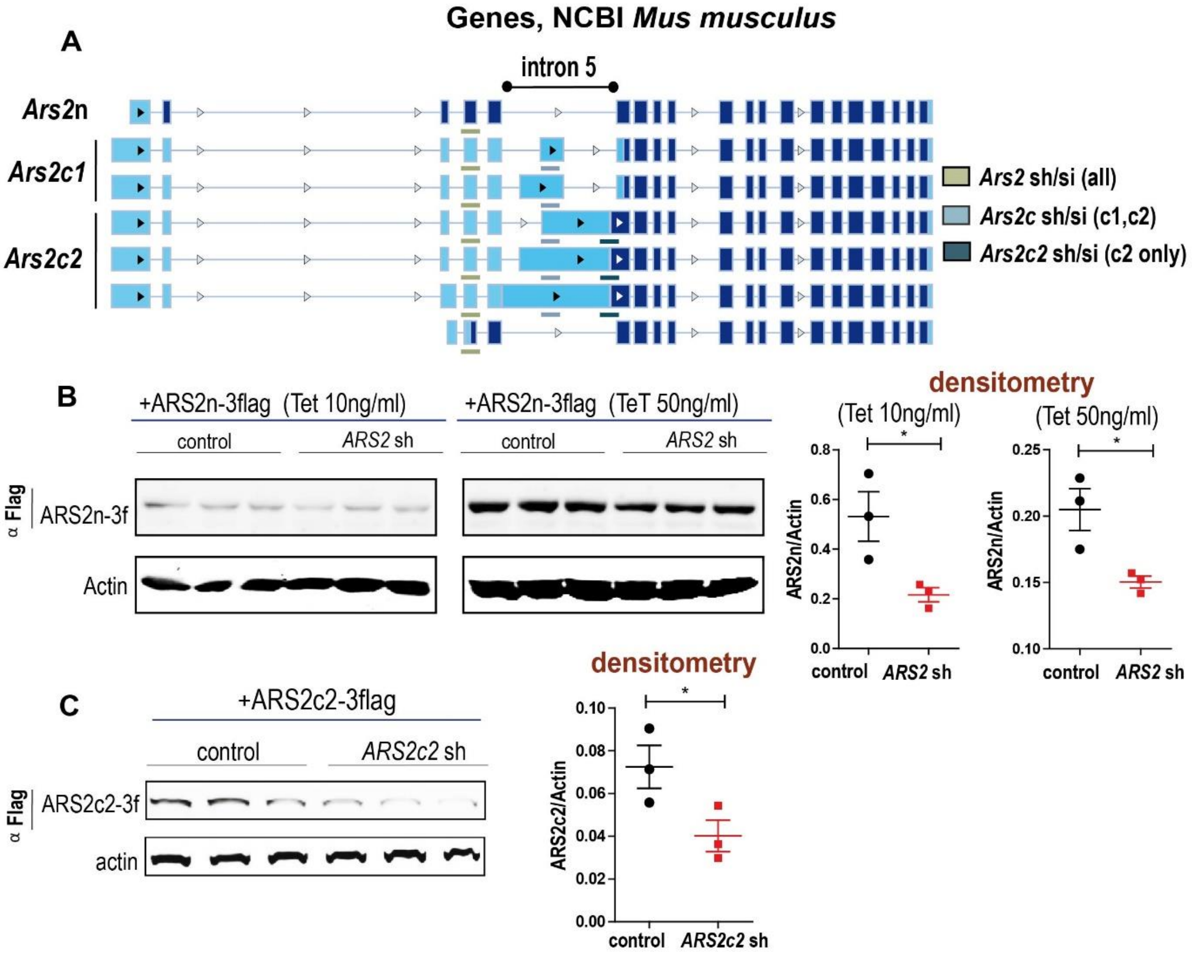


**Sup Figure 4: Validation of shRNAs used in the study. A)** Diagram of shRNAs/RNAi used in the experiments. *Ars2* sh/si targets all *Ars2* isoforms, *Ars2c* sh/si targets all cytoplasmic isoforms and *Ars2c2* sh/si is specific for the *Ars2c2* isoform. **B) C)** Flp-In T-REx 293 inducible cell lines were transfected with *Ars2* sh/*Ars2c2* sh/ control for 48h. After transfection, expression of ARS2n **(B)** or ARS2c2 **(C)** was induced with 10 or 50 ng/ml of Tetracycline for 36h. **B) C)** Flag-tagged ARS2n and ARS2c were detected by Western Blot using anti-flag antibodies. Densitometry is shown in **B)** and **C) (right).** Actin is used as a loading control. Data are represented as mean ± SEM for n=3 biologically independent samples**.** Statistical analysis: One-tail unpaired T-test (*p≤0.05).

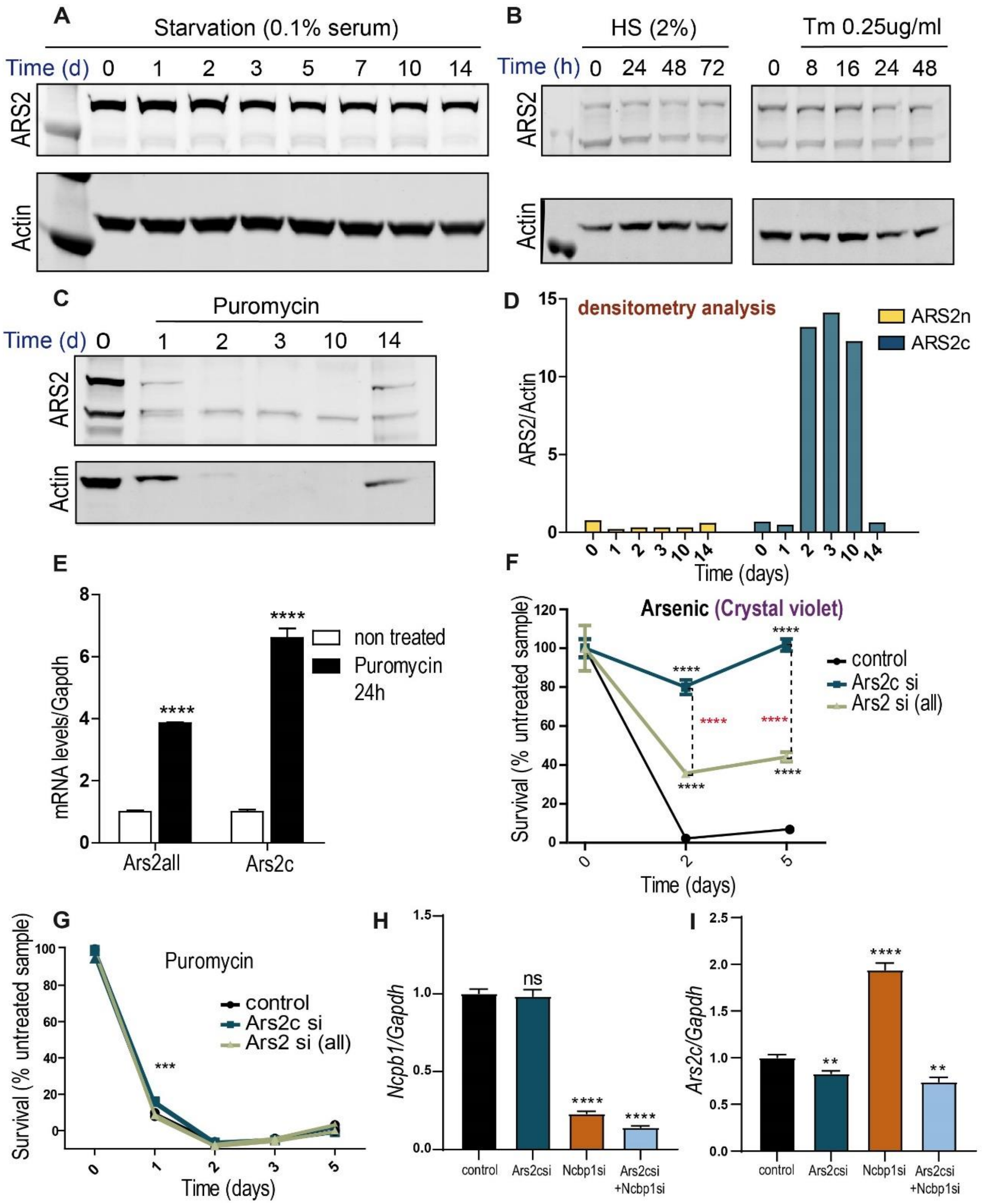


**Sup Figure 5: Expression of ARS2 isoforms is differentially regulated during translation stress.** HeLa **(A, C)** and $C_2C_{12}$ cells **(B, E)** were exposed to different stress conditions: Serum starvation **(A)**, differentiation induction (Horse serum 2%) **(B)**, Tunicamycin **(B)** or Puromycin 2-5 ug/ml **(C)** and **(E)**. ARS2 isoforms were detected by western blot **(A, B, C)**, using an antibody against ARS2, or by RT-qPCR **(E)** using primers to amplify intron 5 (*Ars2c* specific) or all *Ars2* derived transcripts. **C)** Total Protein concentration at time 0 is 30 ug, but due to sample limitation, is 15 ug from times 1-14 days. **D)** Densitometry analysis of **C)**, where actin was used as a normalizer. **E)** Data are represented as mean ± SEM for n=3 biologically independent samples**.** Statistical analysis: two-tail unpaired T-test (****p≤0.0001). **F)** $C_2C_{12}$ cells transfected with *Ars2c*/*Ars2*all/control RNAi were treated with arsenic 40uM **(F)** or puromycin 2 ug/ml **(G)**

for 1-5 days. Survival (% treated/untreated sample) was measured by crystal violet **(F)** or WST1 **(G).** Data are represented as mean ± SEM for n=3 biologically independent samples**.** Statistical analysis: One-way Anova, post-test Tukey's (***p≤0.001, ****p≤0.0001). **H, I)** Validation of Ars2c+Ncbp1si combination. C2C12 cells were transfected with *Ars2c/Ncbp1/Ars2c+Ncbp1*/control RNAi for 48h. *Ncbp1* and *Ars2c* were detected by RT-qPCR. Data are represented as mean ± SEM for n=3 biologically independent samples**.** Statistical analysis: One-way Anova, post-test Dunnett's (**p≤0.01, ****p≤0.0001).

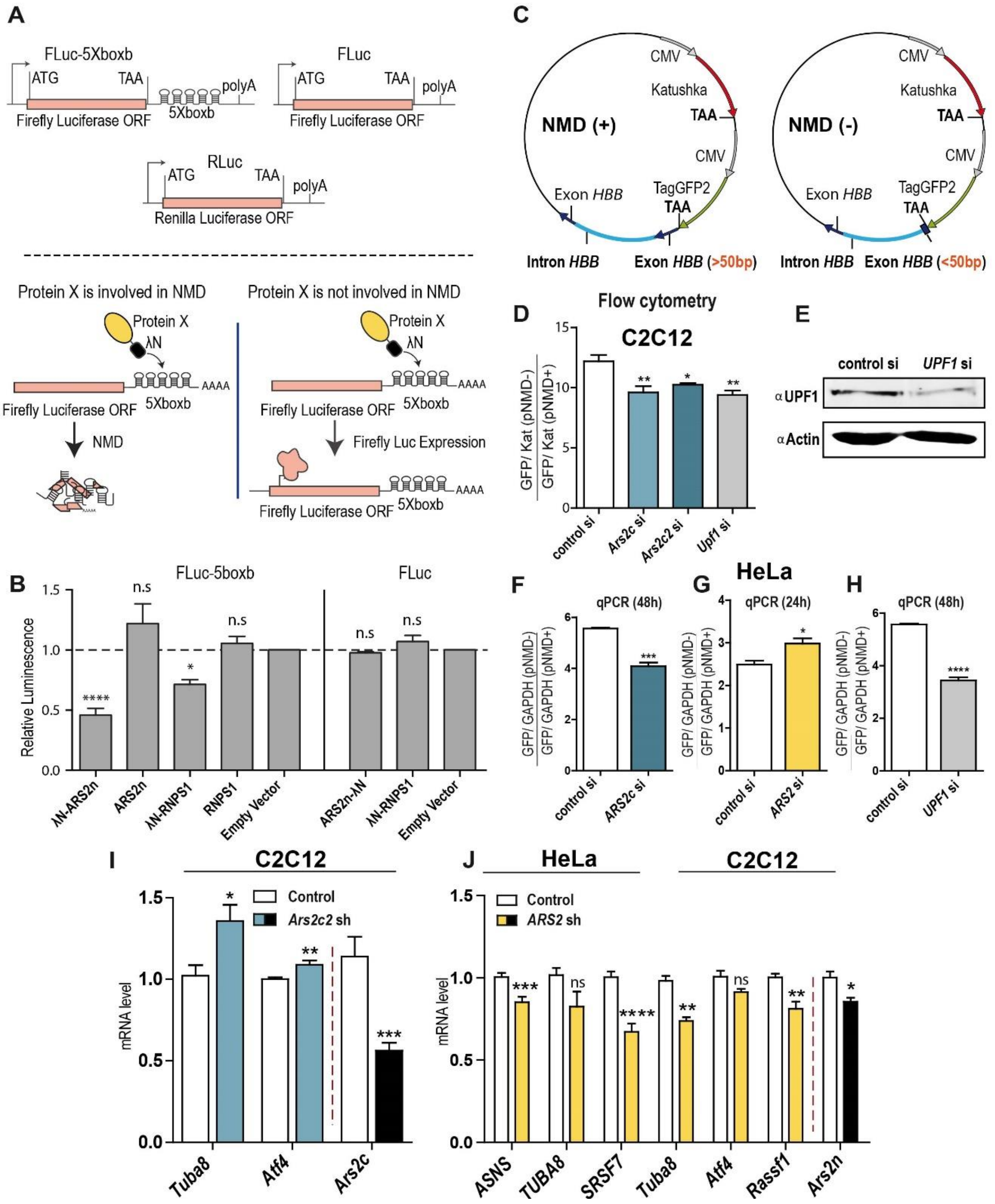


**Sup Figure 6: ARS2 isoforms differentially regulate nonsense mediated decay. A)** Schematic representation of the tethering assay. Protein X-λN fusions are tethered to 5xBoxB sites in the 3'UTR of a

Firefly luciferase reporter. If protein X is involved in NMD, tethering to the reporter results in a loss of Firefly luciferase activity. **B)** HeLa cells were transfected with λN-ARS2n/ ARS2n/ λN-RNPS1/ RNPS1/ control + Fluc-5boxb/Fluc+ Renilla luciferase (normalizer). Luciferase activity was measured at 24h after transfection. Data are represented as mean ± SEM for n=3 biologically independent samples**.** Statistical analysis. One-way Anova, post-test Dunnett's. (*p≤0.05; ****p≤0.0001). Targeting of ARS2n to 3'UTR mediates the degradation of the reporter, similar to NMD (EJC) component RNPS1. **C)** Schematic representation of pNMD+/pNMD- reporters. pNMD+ has an EJC deposited >50 nucleotides from the TagGFP2 stop codon and therefore is degraded by NMD. pNMD- with an EJC placed <50 nucleotides from TagGFP2 stop codon is insensitive to NMD degradation. Both reporters are normalized against the protein Katushka. **D)** $C_2C_{12}$ cells were transfected with pNMD+/pNMD- and *Ars2c*/*Ars2c2*/*Upf1* or control RNAi. Expression levels were detected by Flow cytometry. The pNMD-/pNMD+ ratio is represented in the graphs. Data are represented as mean ± SEM for n=3 biologically independent samples**.** Statistical analysis: One-way Anova, post-test Dunnett's (*p≤0.05; **p≤0.01). **E)** Western blot validation of UPF1 siRNA. Endogenous UPF1 is detected using anti UPF1 antibodies. Actin is used as a normalizer. **F) G) H)** HeLa cells were transfected with pNMD+/pNMD- and *ARS2c*/*ARS2*/*UPF1* or control RNAi. Expression levels were detected by RT-qPCR. The pNMD-/pNMD+ ratio is represented in the graphs. Data are represented as mean ± SEM for n=3 biologically independent samples**.** Statistical analysis: two-tail unpaired T-test (*p≤0.05; ***p≤0.001, ****p≤0.0001). **I) J)** RT-qPCR performed after transfection of HeLa and $C_2C_{12}$ cells with *ARS2* sh/*ARS2c* sh or control for 48h. Each gene was normalized against *GAPDH*. Data are represented as mean ± SEM for n=3 to n=6 biologically independent samples**.** Statistical analysis: two-tail unpaired T-test (*p≤0.05; ** p≤0.01; ***p≤0.001, ****p≤0.0001).

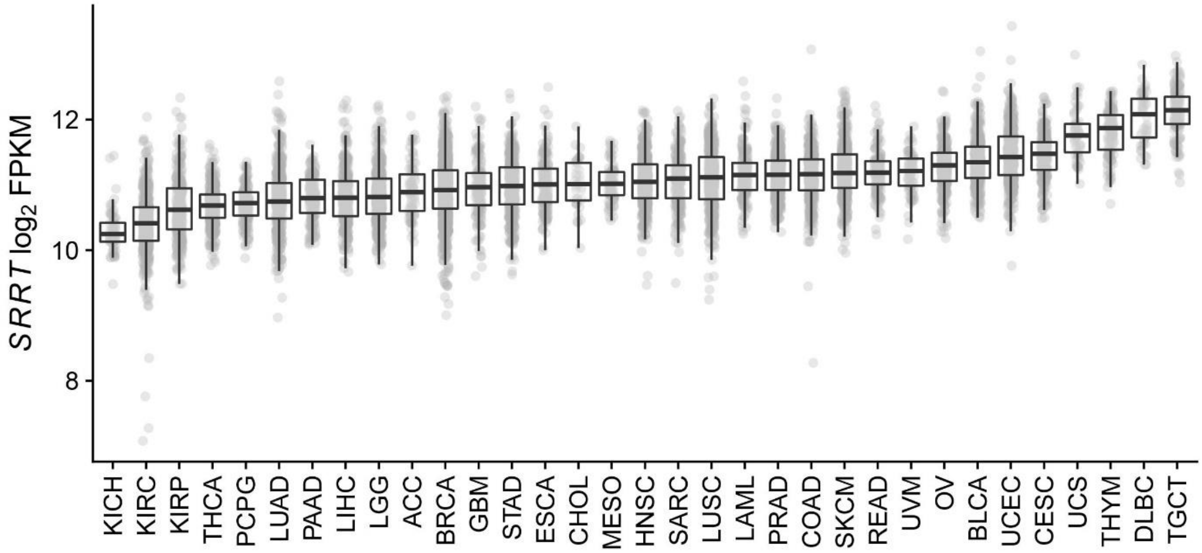


**Figure 7: *ARS2 (SRRT)* expression in cancer.** *ARS2* gene expression for solid cancers available through The Cancer Genome Atlas (TCGA). Each dot represents a patient tumor sample.
